# Structural dynamics of mixed-subunit CaMKIIα/β heterododecamers filmed by high-speed AFM

**DOI:** 10.1101/2025.04.24.650392

**Authors:** Keisuke Matsushima, Takashi Sumikama, Taisei Suzuki, Mizuho Ito, Yutaro Nagasawa, Ayumi Sumino, Holger Flechsig, Tomoki Ogoshi, Kenichi Umeda, Noriyuki Kodera, Hideji Murakoshi, Mikihiro Shibata

## Abstract

CaMKII predominantly assembles into a 12-meric ring assembly, primarily consisting of CaMKIIα and CaMKIIβ variants in the brain. Previous biochemical studies have reported varying ratios of these CaMKII variants across different brain regions and developmental stages. However, direct evidence for the formation of CaMKIIα/β heterooligomers within a 12-meric ring assembly has been lacking at the single-molecule level. Here, we employed high-speed atomic force microscopy to visualize the conformational dynamics of forebrain-mimicked CaMKIIα/β at a 3:1 ratio. Our findings revealed that the CaMKIIα and CaMKIIβ subunits are intermixed within the 12-meric ring assembly, with more than 83% probability that CaMKIIβ subunits adjacent to one another. Furthermore, in the activated state, CaMKIIα/β heterooligomers form a stable kinase domain complex via interactions between adjacent CaMKIIβ subunits, resulting in a long-lasting structure with an exposed target binding site. Collectively, our observations provide insights into the structural role of CaMKIIβ subunits within the CaMKIIα/β heterododecamer.

## Main

CaMKII is a holoenzyme that is abundant in neurons of the brain, particularly in postsynaptic regions, and is essential for the induction of long-term potentiation (LTP) and long-term depression (LTD), both of which serve as cellular models of learning and memory^1–7^. Knockout mice lacking the α variant of CaMKII (CaMKIIα) exhibit deficits in place memory^8^, indicating a direct relationship with memory processes^9^. The CaMKIIβ variant is also present in large amounts in brain neurons, and both CaMKIIα and CaMKIIβ play crucial roles in regulating brain function^7,10,11^. Recent studies have associated perturbations in CaMKIIβ expression with various neuropsychiatric and neurodevelopmental disorders^9,10,12^.

CaMKIIα and CaMKIIβ are composed of a kinase domain, an autoinhibitory-regulatory segment, a linker region, and a hub domain and are activated by Ca^2+^/CaM binding (Fig. 1a and Extended Data Fig. 1)^13–17^. This binding releases the autoinhibitory-regulatory segment from the kinase domain, thereby exposing the active site of the kinase domain and the primary phosphorylation site (Thr286 for CaMKIIα or Thr287 for CaMKIIβ), which facilitates *trans*-autophosphorylation within the oligomer (Extended Data Fig. 1). The subsequent dissociation of Ca^2+^/CaM exposes a second phosphorylation site (Thr305/306 for CaMKIIα or Thr306/307 for CaMKIIβ), enabling additional autophosphorylation. These phosphorylation modifications are reverted to the basal state through dephosphorylation by phosphatases (Extended Data Fig. 1). Although CaMKIIα and CaMKIIβ share an activation mechanism, the primary distinction between them lies in the number of amino acids (a.a.) in the linker region (CaMKIIα: 33 a.a., CaMKIIβ: 96 a.a., Fig. 1 and Extended Data Fig. 2). CaMKIIβ has a longer linker than other variants (γ and δ), and interspecies differences in CaMKII are also concentrated within this linker region^18–20^. Structurally, electron micrographs have revealed that the homooligomers of both CaMKIIα^21^ and CaMKIIβ^22^ in recombinant proteins predominantly form a 12-meric ring assembly. The central hub assembly, which is responsible for organizing each subunit, consists of a hub domain, with the kinase domains distributed around it (Fig. 1b).

**Fig. 1:**
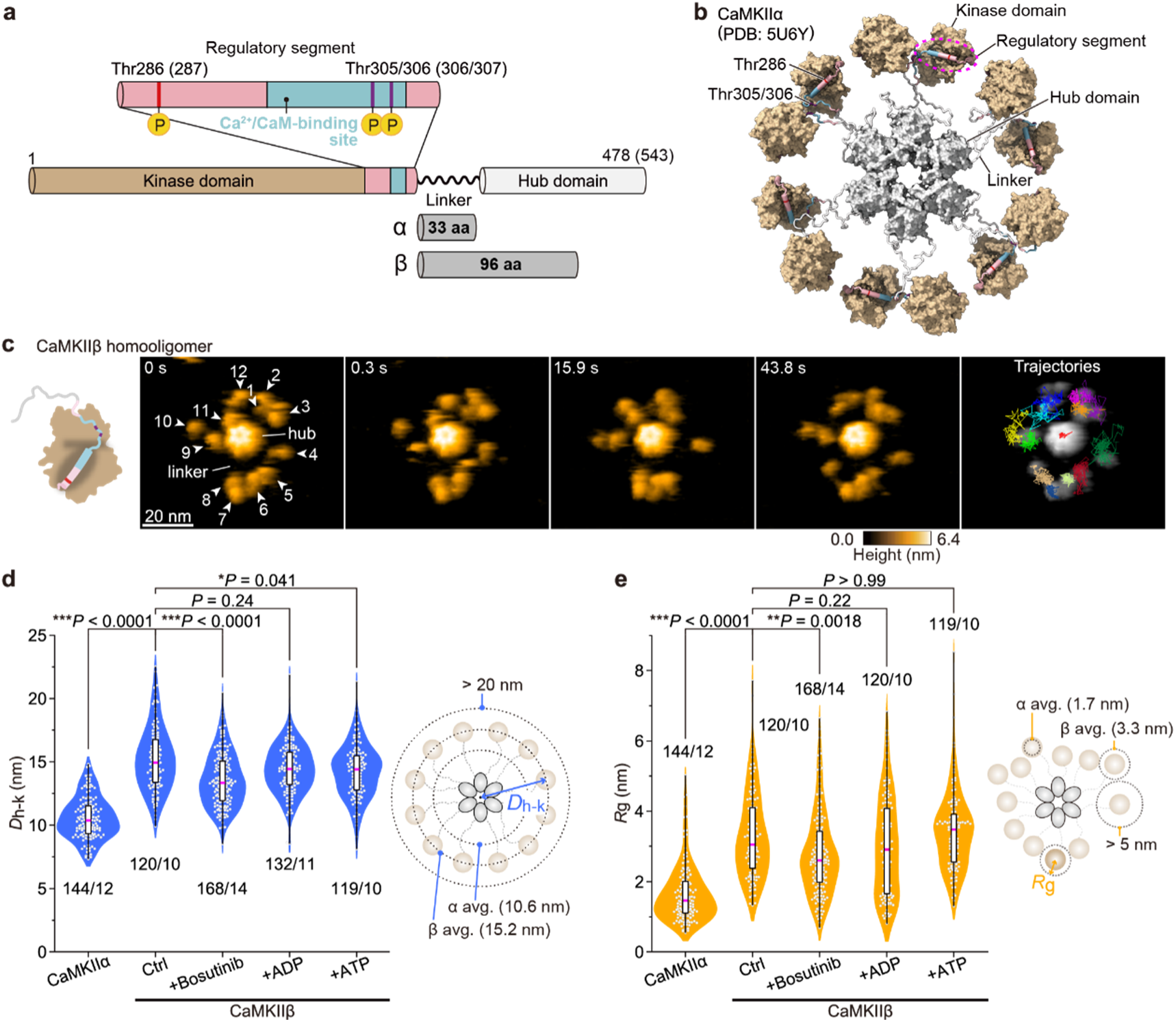
The kinase domains of CaMKIIβ exhibit greater mobility away from the central hub assembly than do those of CaMKIIα. **a**, Domain structure of CaMKIIα and CaMKIIβ. The numbers correspond to the amino acid sequences. The numbers in parentheses indicate CaMKIIβ. **b**, Pseudoatomic model of the 12-meric CaMKIIα from EM data^21^. The solvent-excluded surface is represented. The hub domains (light gray surface, residues 345–472), kinase domains (tan surface, residues 1–273), regulatory segment (magenta cylinder, residues 274–314) with the Ca^2+^/CaM-binding site (blue cylinder, residues 293–310), and linker region (white ribbon, residues 315–344) are shown. The phosphorylation sites at Thr286, Thr305, and Thr306 are highlighted in red and purple. **c**, Sequential HS-AFM images of CaMKIIβ (Ctrl; see also Supplementary Video 1). The white arrowheads indicate kinase domains with arbitrary numbering. Frame rate, 3.3 frames/s. The trajectories of the kinase domains and the center of the hub assembly (red in the center) were tracked for approximately 45 s. **d**,**e**, *D*_h-k_ (**d**) and *R*_g_ (**e**) under the indicated conditions. The number of samples (kinases/holoenzymes) is indicated in the figure. Right: Illustrations of *D*_h-k_ and *R*_g_. Comparisons between CaMKIIα and CaMKIIβ (Ctrl) (Mann–Whitney U test) and CaMKIIβ under different conditions (Kruskal‒Wallis test with Dunn’s post hoc test).

Interestingly, previous biochemical analyses have revealed that native CaMKII in neurons has varying ratios of variants across different brain regions. For example, in the adult forebrain, the α:β ratio is 3:1, whereas in the cerebellum, this ratio is 1:4 (ref.^23–27^). Varying variant ratios have also been observed during development, with a 1:1 ratio reported in the forebrain during the first 10 days of mouse development^28,29^. However, whether these ratios arise from heterooligomers in a CaMKII 12-mer or reflect different ratios of homooligomers of CaMKII remains a topic of debate.

How does the composition of the 12-meric ring assembly of CaMKII, as a heterooligomer or homooligomer, influence its biological function in neurons? Previous studies have indicated that only CaMKIIβ binds to F-actin via its linker region^30^, playing a crucial role in its localization within neurons^31–33^. If the CaMKIIα/β heterooligomer contains a smaller number of CaMKIIβ subunits, as observed in the forebrain, the binding of the CaMKIIα/β heterooligomer to F-actin is expected to be relatively weak, since only three CaMKIIβ subunits in the 12-mer can bind. Thus, the localization of the CaMKIIα/β heterooligomer to F-actin appears to require an additional mechanism. In contrast, CaMKII in the cerebellum, which is characterized by a higher ratio of CaMKIIβ subunits and fewer CaMKIIα subunits, can bind strongly to F-actin because 9 to 10 CaMKIIβ subunits in the 12-mer able to bind to F-actin. If CaMKII forms homooligomers, the colocalization of CaMKIIα and CaMKIIβ with F-actin would not occur, and each variant exists independently within neurons, with only CaMKIIβ binding to F-actin. In the context of structural plasticity in the spine, the stimulation of LTP and LTD is thought to induce the binding and dissociation of CaMKII from F-actin, leading to substantial changes in spine volume^34,35^. Therefore, obtaining direct evidence of the subunit composition of the 12-meric ring assembly of CaMKII could clarify its binding mode with F-actin and greatly improve our understanding of the molecular mechanism underlying synaptic plasticity. However, thus far, biochemical experiments have only indirectly verified the stoichiometry of CaMKII variants within CaMKII oligomers through electrophoresis^24^, leaving the question of hetero-versus homooligomer composition open to speculation.

High-speed atomic force microscopy (HS-AFM) is a unique technique that allows direct visualization of the nanometer-scale dynamics of functional proteins^36,37^, protein‒DNA complexes^38^, and individual subunits within oligomers^39^ in a liquid environment.

Previously, we applied HS-AFM to homooligomers overexpressing CaMKIIα, enabling visualization of the activation state-dependent structural dynamics of the kinase domain at the sub-molecular level^39^. In this study, we employed HS-AFM to directly visualize the dynamics of individual subunits within a CaMKIIα/β heterooligomer with a 3:1 α:β ratio, as well as a CaMKIIβ homooligomer within a 12-meric ring assembly. Our findings provide insights into the dynamics of individual subunits in CaMKII 12-meric ring assembly and suggest biological roles on the basis of these dynamics.

## Results

### The kinase domains of CaMKIIβ exhibit greater mobility away from the central hub assembly than do those of CaMKIIα

We previously characterized the CaMKIIα subunit in homooligomers by measuring the distance (*D_h-k_*) from the central hub assembly to each kinase domain and the mobility (*R_g_* in 45 seconds) of each kinase domain^39^. Since CaMKIIβ possesses a longer linker than CaMKIIα does, it is possible to distinguish the CaMKIIα and CaMKIIβ subunits in the heterooligomer by comparing *D_h-k_* and *R_g_* values. Therefore, we initially applied HS-AFM to the CaMKIIβ homooligomer to characterize the dynamics of the CaMKIIβ subunits. To facilitate single-molecule observations of CaMKIIβ, we overexpressed CaMKIIβ in HEK293 cells and purified it using two different tags, similar to previous HS-AFM studies of CaMKIIα (Extended Data Fig. 3a)^39^. The purified CaMKIIβ was then immobilized on a positively charged pillar–arene substrate^39,40^ to enable visualization of the dynamics of the kinase domains. The resulting HS-AFM images of the CaMKIIβ homooligomers revealed a gear-like structure with six protrusions at the center surrounded by scattered spherical domains (Fig. 1c and Supplementary Video 1). Compared with previous electron micrographs of CaMKIIβ , the shape observed in this structure allows us to attribute the central gear-like structure to the hub assembly, whereas the surrounding domains correspond to the kinase domains. To quantify the mobility of the kinase domains of the CaMKIIβ subunits, we calculated *D_h-k_* and *R_g_* on the basis of the coordinates of the center of the hub assembly and the maximum height of each kinase domain (see the Methods). We found that the kinase domain of CaMKIIβ in the basal state presented an average *D_h-k_* of 15.2 ± 2.5 nm and an *R_g_* of 3.3 ± 1.2 nm (Fig. 1d, e). The *D_h-k_*values were consistent with those of previously reported electron micrographs of CaMKIIβ^22^. Furthermore, the *D_h-k_* and *R_g_*values were significantly greater than observed in our HS-AFM results for CaMKIIα (Fig. 1d, e). These findings suggest that the length of the linker (α: 33 a.a., β: 96 a.a.) plays a major role in determining the positioning and mobility of the kinase domains. Additionally, HS-AFM observations in the presence of bosutinib, an ATP-competitive inhibitor of CaMKII, revealed significant reductions in both *D_h-k_*and *R_g_* values (Fig. 1d, e, Extended Data Fig. 4 and Supplementary Video 2). This finding suggests that, in addition to linker length, the interaction between the regulatory segment and the kinase domain is also important for kinase domain mobility (a stronger interaction of the regulatory segment correlates with decreased mobility of the kinase domains). In the presence of ADP or ATP, *R_g_* did not significantly differ from that in the control experiment. These findings indicate that bosutinib, which has a stronger binding affinity than nucleotides do, enhances binding to the active site of the regulatory segment, resulting in reduced kinase domain mobility (Fig. 1d, e). Thus, the inhibitory effect of bosutinib is consistent across CaMKIIα and CaMKIIβ; however, the positioning relative to the central hub assembly and the mobility of the kinase domains differ between CaMKIIα and CaMKIIβ. This distinction enables us to distinguish between the two subunits of the CaMKIIα/β heterooligomer.

### The CaMKII 12-meric ring assembly comprises a mixture of CaMKIIα and CaMKIIβ subunits, with the CaMKIIβ subunits positioned adjacently

In forebrain neurons, biochemical studies have reported a ratio of CaMKIIα and CaMKIIβ of 3:1 (ref.^23,24^). Accordingly, we prepared purified proteins with different CaMKII variant ratios via overexpression in HEK293 cells at an α:β ratio of 3:1 (see the Methods; c) and subsequently performed HS-AFM. HS-AFM images revealed a 12-meric ring assembly resembling the structures observed for the CaMKIIα and CaMKIIβ homooligomers (Fig. 2 and Supplementary Video 3). We analyzed the *D_h-k_* and *R_g_* values for each kinase domain in the CaMKIIα/β heterooligomers to identify which kinase domains in the 12-meric ring assembly corresponded to the CaMKIIα and CaMKIIβ subunits. The identification criteria are shown on the right side of Figure 2C (see the Methods for details). Notably, distinguishing CaMKIIβ subunits solely on the basis of *D_h-k_* values was challenging. The mobility of the kinase domains, as indicated by *R_g_*, was critical for accurately identifying the CaMKIIβ subunit. The results indicated that CaMKIIα and CaMKIIβ form heterooligomers intermixed at a ratio of approximately 3:1 within the 12-meric ring assembly (Fig. 2c). This ratio showed considerable variation (Fig. 2c and Extended Data Fig. 5a). Interestingly, when two or more CaMKIIβ subunits were present in the 12-meric ring assembly, they were more likely to be adjacent rather than randomly positioned, with probabilities of approximately 45% (18.2% for random positioning) and 100% (49.1% for random positioning) for two and three CaMKIIβ subunits, respectively, resulting in an overall probability of 84% (*n* = 37 oligomers)

**Fig. 2:**
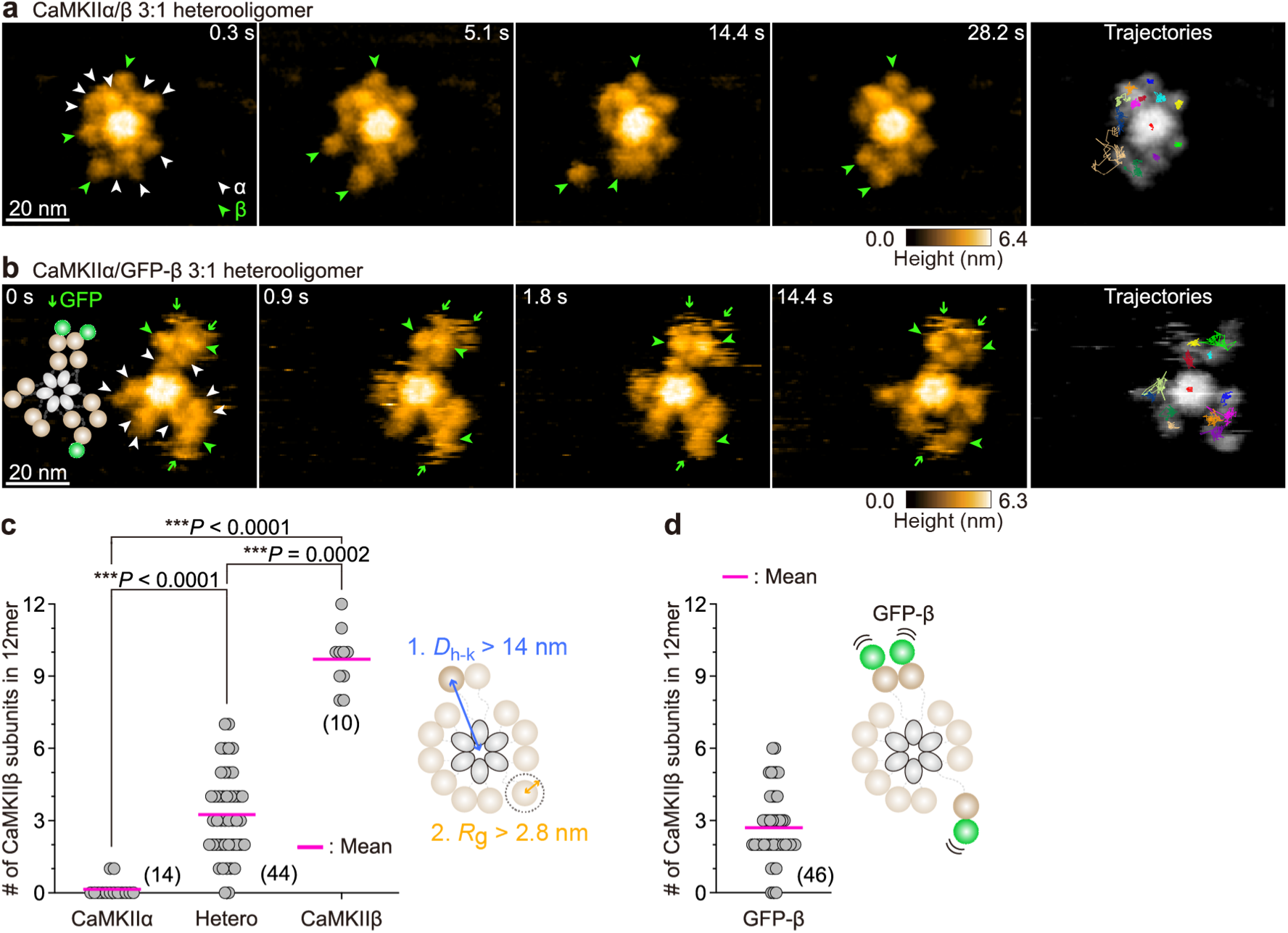
The CaMKII 12-meric ring assembly consists of a mixture of CaMKIIα and CaMKIIβ subunits, with CaMKIIβ subunits positioned adjacent to each other. **a**,**b** Sequential HS-AFM images of the CaMKIIα/β 3:1 heterooligomer (**a**; see also Supplementary Video 3) and the CaMKIIα/GFP-β 3:1 heterooligomer (**b**; see also Supplementary Video 4). The white and green arrowheads indicate the kinase domains of CaMKIIα and CaMKIIβ, respectively. The green arrows denote GFP (**b**). Frame rate, 3.3 frames/s. The trajectories of the kinase domains and the center of the hub assembly (red in the center) were tracked for approximately 30 s. **c**,**d**, The number of kinase domains attributed to CaMKIIβ subunits in the 12-meric ring assembly. To test the validity of this criterion, we applied the same criterion to the CaMKIIα homooligomer and CaMKIIβ homooligomer and assessed the number of molecules classified as CaMKIIβ subunits. The number in parentheses indicates the analyzed CaMKII oligomers (Kruskal‒Wallis test with Dunn’s post hoc test, **c**).

To further confirm the ratio and localization of the CaMKIIβ subunit in the 12-meric ring assembly, we prepared GFP-CaMKIIβ by adding GFP to the N-terminus of the CaMKIIβ subunit and subsequently purified a CaMKIIα:GFP-CaMKIIβ heterooligomer at a ratio of 3:1 (see the Methods; Extended Data Fig. 3). In this case, the CaMKIIβ subunit was labeled with GFP (small sphere), allowing for visual recognition by HS-AFM. Under our AFM experimental conditions, GFP did not strongly adsorb to the AFM substrate and appeared as a highly flexible protrusion (green arrow in 4). As expected, the CaMKII heterooligomer included the GFP-fused kinase domain (CaMKIIβ subunit), facilitating the determination of the α:β ratio and localization. Similar to the conditions without GFP, the CaMKIIα and GFP-CaMKIIβ subunits were intermixed in the 12-meric ring assembly at a ratio of approximately 3:1 (Fig. 2d), and the probability of adjacent GFP-CaMKIIβ subunits was 78% (*n* = 40 oligomers) (Extended Data Fig. 5b,c). Previously, the presence of heterooligomers in neurons has been inferred from differences in molecular weight assessed via electrophoresis^24^, but direct evidence of their existence has not been reported. Single-molecule imaging using HS-AFM demonstrated that CaMKII forms a heterooligomer with intermixed CaMKIIα and CaMKIIβ subunits, with CaMKIIβ subunits positioned adjacently within a 12-meric ring assembly.

### The kinase domains of CaMKIIβ in the Ca^2+^/CaM-bound state exhibit low mobility away from the central hub assembly and form a stable kinase domain complex

Next, we aimed to elucidate the effect of the adjacent CaMKIIβ subunit in the 12-meric ring assembly on the active state of CaMKII. Initially, we premixed the CaMKIIβ homooligomer with Ca^2+^/CaM in tubes and performed HS-AFM to characterize the binding state of Ca^2+^/CaM in the CaMKIIβ subunit clearly. An additional globular structure was observed near the kinase domain (Fig. 3a and Supplementary Video 5). Given that the Ca^2+^/CaM binding site of CaMKIIβ is located close to the kinase domain (Fig. 1a) and that these globular structures dissociated during HS-AFM scanning, we concluded that they correspond to Ca^2+^/CaM bound to the CaMKIIβ subunits. The *D_h-k_* value of the Ca^2+^/CaM-bound kinase domain was 17.9 ± 2.7 nm, which was 2.7 nm greater than that of the control, whereas the *R_g_* value was 2.1 ± 1.2 nm, which was significantly lower than that of the control (Fig. 3d, e). Similarly, in the CaMKIIα/β heterooligomer, the Ca^2+^/CaM-bound kinase domain exhibited an extended structure (Extended Data Fig. 6a and Supplementary Video 6). Our previous HS-AFM results for CaMKIIα homooligomers also revealed an extended structure of the kinase domain owing to Ca^2+^/CaM binding, with a length of approximately 3 nm (ref.^39^). Although CaMKIIβ contains approximately three times more amino acid residues in the linker region than does CaMKIIα, the mechanism by which the regulatory segment is released upon Ca^2+^/CaM binding to the kinase domain (activation mechanism) is consistent for both CaMKIIα and CaMKIIβ.

**Fig. 3:**
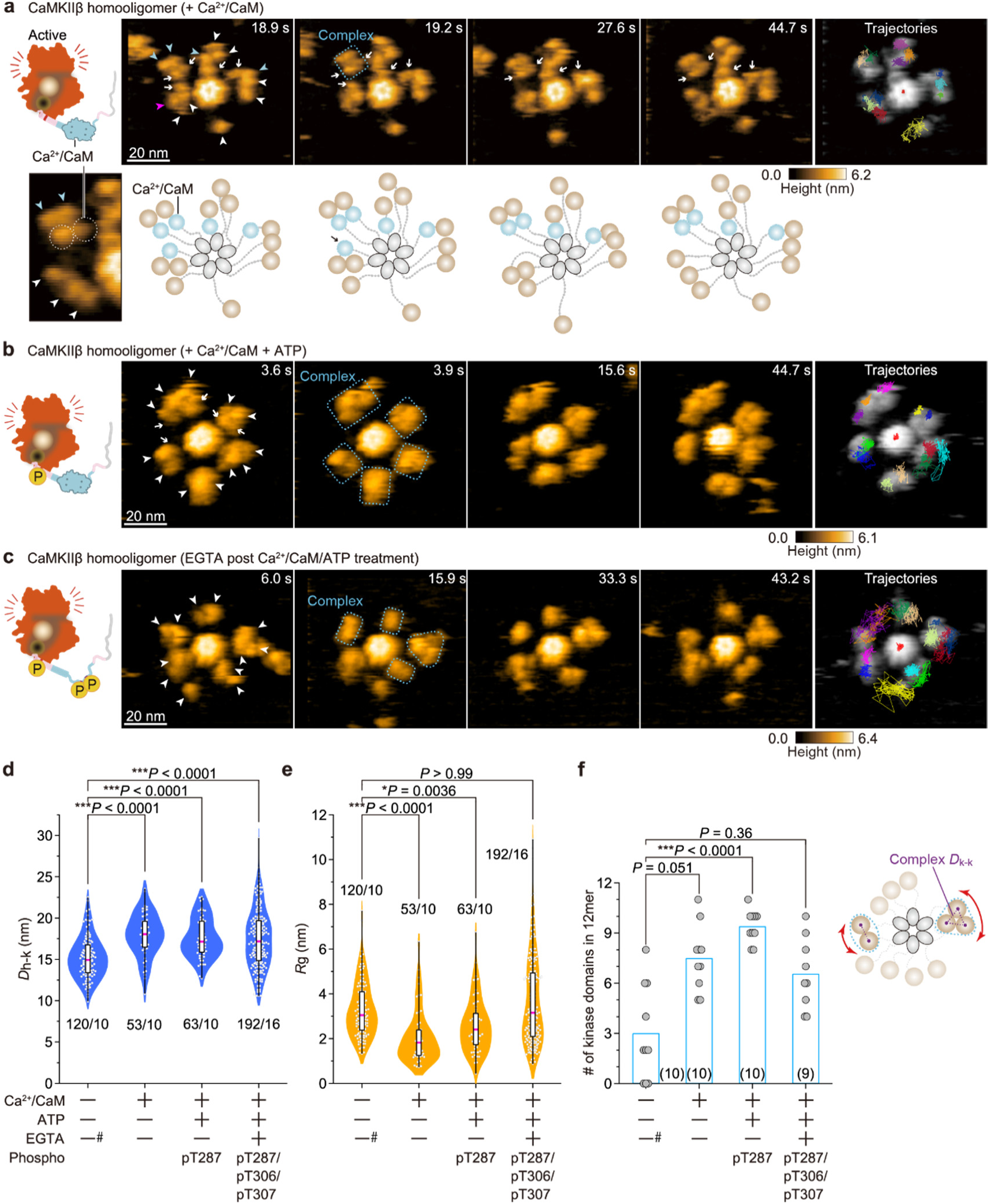
The kinase domains of CaMKIIβ in the activated state form a stable kinase domain complex. **a-c**, Sequential HS-AFM images of Ca²⁺/CaM-bound CaMKIIβ (**a**; 1 mM Ca^2+^, 800 nM CaM; see also Supplementary Video 5), pT287 CaMKIIβ (**b**; 1 mM Ca^2+^, 800 nM CaM, and 1 mM ATP; see also Supplementary Video 7), and pT287/pT306/pT307 CaMKIIβ (**c**; see also Supplementary Video 9). CaMKIIβ was first activated to induce pT287, as described previously (Extended Data Fig. 1). Subsequently, EGTA (2 mM) was added to induce Ca^2+^/CaM dissociation and pT306/pT307 (autophosphorylation) (**c**). The white arrows indicate Ca^2+^/CaM bound to the kinase domains. The white, blue and magenta arrowheads indicate the kinase domains, Ca²⁺/CaM binding and dissociating kinase domains, respectively. The blue dotted squares indicate a kinase domain complex. Diagrams of the presumed structures of the linker regions in 12-meric CaMKIIβ are also shown at the bottom of each HS-AFM image (**a**). Frame rate, 3.3 frames/s. The trajectories of the kinase domains and the center of the hub assembly (red in the center) were tracked for approximately 45 s. **d**,**e**, *D*_h-k_ (**d**) and *R*_g_ (**e**) under the indicated conditions. The number of samples (kinases/holoenzymes) is indicated in the figure (Kruskal‒Wallis test with Dunn’s post hoc test). #: 0.1 mM EGTA **f**, The number of kinase domains that remained in the complex state for more than 45 s per 12-meric CaMKIIβ homooligomer under the indicated conditions. The number in parentheses is the number of holoenzymes used in the analysis (Kruskal‒Wallis test with Dunn’s post hoc test). Right: Illustrations of Complex *D*_k-k_.

Interestingly, we found that the binding of Ca^2+^/CaM increases the number of kinase domains that move together in close proximity for extended periods (>45 s during HS-AFM observation) (blue box at 19.2 s in Fig. 3a). We defined this phenomenon as a kinase domain complex (see the Methods). This complex was also observed in the basal state (Extended Data Fig. 7a); however, the binding of Ca^2+^/CaM enables this complex to persist even longer (Fig. 3f). While this kinase domain complex is present in the CaMKIIα subunit, it is more characteristic of the CaMKIIβ subunit, as it forms at a higher frequency in the CaMKIIβ subunit (Extended Data Fig. 7b, c).

### The kinase domains of CaMKIIβ in the Ca^2+^/CaM-bound pT287 state form a more stable kinase domain complex

After CaMKII binds to Ca^2+^/CaM, the first autophosphorylation occurs at the CaMKIIα:Thr286 and CaMKIIβ:Thr287 sites via ATP hydrolysis (Ca^2+^/CaM+pT286 [pT287 in CaMKIIβ], Extended Data Fig. 1). To visualize the structural dynamics of this phosphorylation state, we performed HS-AFM following the premixing of Ca^2+^/CaM and ATP in a tube. The phosphorylation states of both CaMKII variants under these experimental conditions were confirmed by biochemical experiments (Extended Data Fig. 8).

HS-AFM images of the CaMKIIβ homooligomer in the presence of Ca^2+^/CaM and ATP captured Ca^2+^/CaM bound to the proximal site of the kinase domain, as observed without ATP. The *D_h-k_* value was 17.6 ± 2.4 nm, indicating an extension of 2.4 nm compared with that of the control, whereas the *R_g_*was 2.6 ± 1.2 nm, which was significantly lower than that observed in the control (Fig. 3b, d, e, and Supplementary Video 7). Our previous results for CaMKIIα homooligomers demonstrated that Ca^2+^/CaM+pT286 adopts an extended structure of approximately 3 nm. However, while the *R_g_* value was comparable to that in the basal state for CaMKIIα, it remained low for CaMKIIβ.

Furthermore, the number of kinase domains that maintained the complex state for a long time significantly increased in the presence of Ca^2+^/CaM and ATP (blue boxes at 3.9 s in Fig. 3b and Fig. 3f). This observation suggests that Ca^2+^/CaM+pT287 of CaMKIIβ forms a more stable kinase domain complex. This complex structure is likely connected to the central hub assembly through multiple CaMKIIβ subunits, which are anchored at several points, resulting in reduced mobility (lower *R_g_* values).

We also observed the formation of kinase domain complexes between adjacent CaMKIIβ subunits in the 12-meric ring assembly of the CaMKIIα/β heterooligomers (Extended Data Fig. 6b). Most of these complexes were stable and persisted for an extended period (>45 s) (90.2%, *n* = 51 complexes) (Supplementary Video 8). The occurrence of kinase domain complexes between CaMKIIβ subunits in the heterooligomers suggests that the kinase domain of the CaMKIIβ subunit plays a role in stabilizing the active conformation over time.

### The kinase domains of CaMKIIβ in the fully phosphorylated state, like those of CaMKIIα, coexist with a kinase domain with increased mobility and an oligomerized kinase domain

We next compared the structural dynamics of the kinase domains in further phosphorylated states (CaMKIIβ: pT287/pT306/pT307, Extended Data Fig. 1). After the CaMKIIβ homooligomer was incubated with Ca^2+^/CaM and ATP in a tube at 30°C for 5 min, we added EGTA to dissociate Ca^2+^/CaM and subsequently performed HS-AFM. The phosphorylation state of CaMKIIβ under these experimental conditions was confirmed by biochemical experiments (Extended Data Fig. 8).

The *D_h-k_* value of the fully phosphorylated state of the CaMKIIβ homooligomer was 17.5 ± 3.5 nm, which indicated significant extension (by 2.3 nm) compared to that observed in the control experiments. The *R_g_* value was 3.7 ± 2.0 nm, indicating a wide distribution (Fig. 3c, d, e). HS-AFM videos demonstrated that some CaMKIIβ subunits moved far from the central hub assembly and exhibited extensive movement, whereas other subunits showed low mobility, as multiple kinase domains assembled together (Fig. 3c, Extended Data Fig. 6d, and Supplementary Video 9). Similar conformational changes in kinase domains were observed in CaMKIIα homooligomers, characterized by increased mobility and kinase domain oligomerization (KD-oligomerization)^39^. Thus, in the fully phosphorylated state, the conformational changes in individual CaMKIIα and CaMKIIβ subunits are similar. It is likely that phosphorylation at multiple sites disrupts the structure of the dissociated regulatory segment, resulting in increased flexibility. However, when multiple kinase domains are in close proximity due to high flexibility, some kinase domains may undergo conformational changes that lead to the entanglement of disordered regions, resulting in KD-oligomerization and decreased mobility. Stable kinase domain complexes were also observed, although at a lower frequency than in the Ca^2+^/CaM-bound pT287 state (blue boxes at 15.9 s in Fig. 3c and Fig. 3f).

HS-AFM of the fully phosphorylated state in the CaMKIIα/β heterooligomer revealed similar structural changes in both the CaMKIIα and CaMKIIβ subunits (Extended Data Fig. 6c and Supplementary Video 10). Based on these results, the differences observed between the CaMKIIα and CaMKIIβ subunits in the activated state can be attributed to the formation of a stable kinase domain complex among the CaMKIIβ subunits, facilitated by Ca^2+^/CaM binding. Interestingly, in the CaMKIIα/β heterooligomers, stable kinase domain complexes between CaMKIIβ subunits frequently formed because the CaMKIIβ subunits were initially positioned adjacent to each other in the 12-meric ring assembly.

### Adjacent CaMKIIβ subunits in the 12-meric ring assembly decrease kinase activity but do not affect dephosphorylation resistance

How does the kinase domain complex affect CaMKII function? To address this question, we performed biochemical experiments to assess the kinase activity of both CaMKII variants and their resistance to phosphatase-mediated dephosphorylation. Kinase activity was measured on the basis of the phosphorylation of the substrate Syntide-2 (see the Methods). Interestingly, in Ca^2+^/CaM-bound pT286 (pT287 in CaMKIIβ), kinase activity was significantly lower for CaMKIIβ homooligomers, even though the phosphorylation level of Thr286 (Thr287 in CaMKIIβ) was comparable across CaMKII variants and heterooligomers (Fig. 4a-c). In addition, the resistance to dephosphorylation by a phosphatase (PP2A) was similar for CaMKIIα and CaMKIIβ homooligomers in the Ca^2+^/CaM-bound pT286/7 state (Fig. 4d-f). Together with the HS-AFM findings regarding kinase domain complex formation in CaMKIIβ subunits in the activated state, these biochemical analyses indicate that the kinase domain complex likely adopts a structure that reduces kinase activity without affecting dephosphorylation resistance at Thr287.

**Fig. 4:**
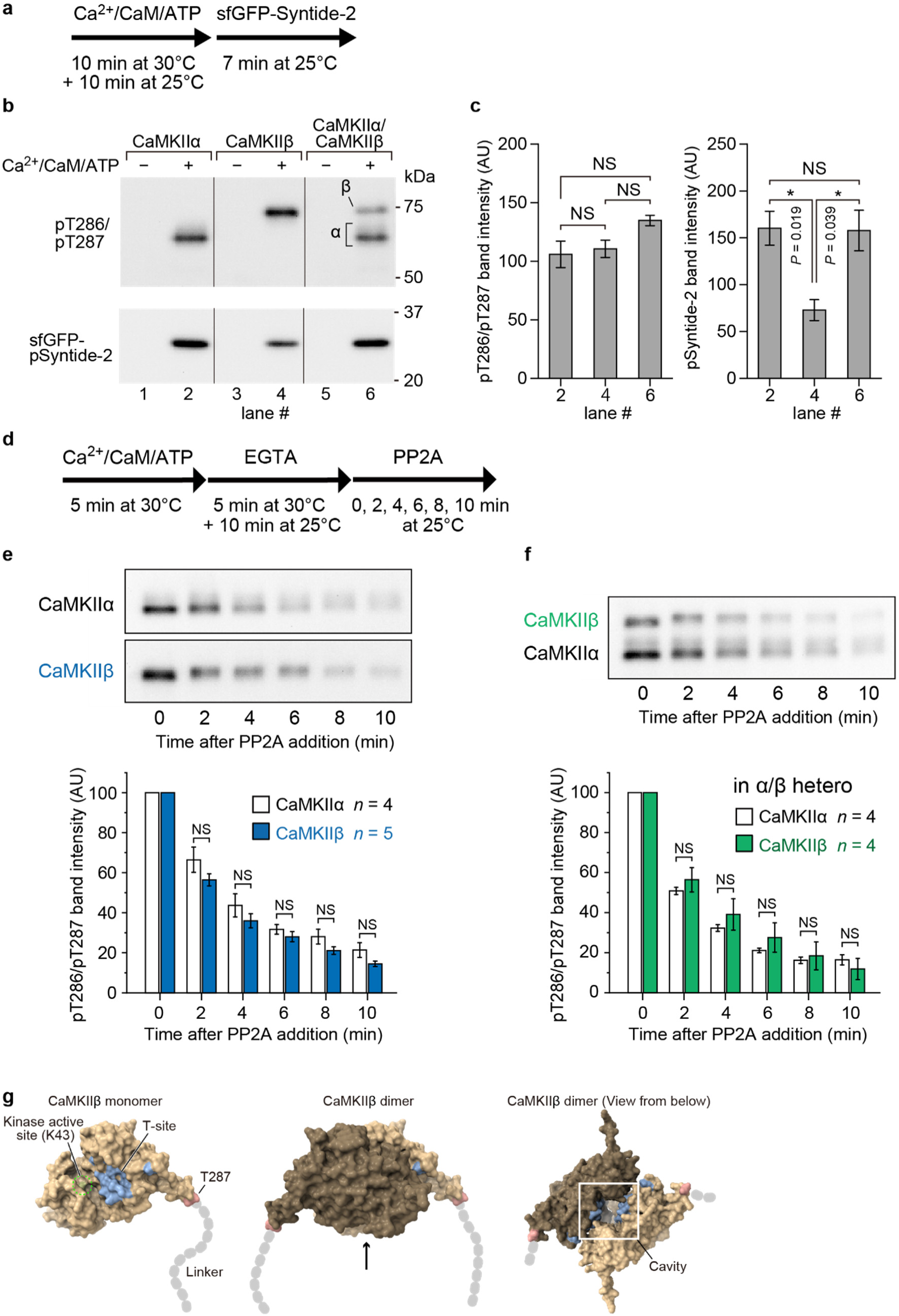
Adjacent CaMKIIβ subunits reduce kinase activity but do not affect dephosphorylation resistance. **a**, Time course of the substrate phosphorylation experiment. **b**, The phosphorylation of sfGFP-syntide-2 by CaMKII holoenzymes was detected by western blotting. CaMKIIα (lanes 1 and 2), CaMKIIβ (lanes 3 and 4), and CaMKIIα/β (lanes 5 and 6) were prepared as described in the Methods. sfGFP-Syntide-2 (1 μM) was incubated for 7 min at 25°C as a substrate for CaMKII. Note that this protocol induces kinase domain complexes in CaMKIIβ holoenzymes (lanes 4 and 6, Fig. 3). **c**, Quantification of the experimental data. The error bars indicate the SEM for four independent experiments. NS (not significant, *P* > 0.05); * *P* < 0.05; ** *P* < 0.01. Following one-way ANOVA, the Bonferroni correction post hoc test was performed. **d**, The time course of the dephosphorylation experiment. **e**,**f**, Representative results of western blotting and quantification of the dephosphorylation of CaMKII holoenzymes by PP2A. pT286 (for CaMKIIα) or pT287 (for CaMKIIβ) CaMKII was prepared as described in the Methods section, and PP2A was added at 0 min. In the case of the rat CaMKII bands, slight smearing occurred, so the dense bottom band was measured. The error bars indicate the SEM, and the number of experiments is shown in the figures. NS (not significant, *P* > 0.05); t test. **g**, Monomeric and dimeric structures of the kinase domain of CaMKIIβ predicted by AlphaFold3. To predict the structure in which the autoinhibitory domain is displaced, we predicted the sequence of the kinase domain at residues 1-287. The dimer structure masks the active site (K43) of the kinase domain. However, owing to the cavity (white box), the T-site surface remains accessible from the direction indicated by the arrow. The solvent-excluded surface is depicted.

We subsequently aimed to elucidate this possible structure by predicting the dimeric configuration of the kinase domain of the CaMKIIβ subunit in the activated state using AlphaFold3 (AF3)^41^. For this prediction, a truncated CaMKIIβ subunit [the autoinhibitory segments and the hub domain were removed] (1-287 a.a.) was used. The predicted dimeric structure revealed that one part of the kinase domain covers the active site of the other kinase domain (Fig. 4g). However, the pT287 surface was found to be free, adopting a conformation accessible to phosphatases. Therefore, the kinase domain complex predicted by AF3 can be interpreted as reducing kinase activity, as the active sites of the kinase domains mask each other, whereas the phosphorylation site (Thr287) remains exposed.

### Kinase domain complexes formed in the activated state exhibit slower structural decay to the basal state after Ca^2+^/CaM dissociation

An additional potential role of the kinase domain complex may involve structural stabilization of the extended structure of the kinase domain that occurs upon Ca^2+^/CaM binding. This structure exposes the active site, as well as the substrate binding site (S-site) and target binding site (T-site), for interactions with other proteins within the kinase domain (Extended Data Fig. 1). To verify the structural role of the kinase domain complex, we compared the decay time of the extended structure of the kinase domain to the basal state between CaMKIIα and CaMKIIβ. Here, we used CaMKIIα_T286A_ and CaMKIIβ_T287A_ mutants, which lack the first phosphorylation site, to demonstrate that the decay of the extended structure is independent of phosphorylation. Specifically, each CaMKIIα_T286A_ and CaMKIIβ_T287A_ homooligomer was mixed with Ca^2+^/CaM in a tube and incubated at 30°C for 5 min to induce the Ca^2+^/CaM binding state. HS-AFM was subsequently performed to observe spontaneous Ca^2+^/CaM dissociation. The decay was calculated by measuring the time from the frame in which Ca^2+^/CaM dissociated until the frame in which the kinase domain returned to its basal state (see the Methods; Fig. 5, Extended Data Fig. 9 and Supplementary Video 11). The results showed that the number of kinase domains that did not return to the basal state during the HS-AFM observation time (>30 s) increased in CaMKIIβ_T287A_ (α: 12.2% [*n* = 41], β: 57.5% [*n* = 45] in Fig. 5g), and most of these domains had formed kinase domain complexes. This finding indicates that the kinase domain complex helps to maintain the extended structure (activated state) of the kinase domain for an extended period. Previous studies have reported that exposure of the S- and T-sites facilitates interactions with a variety of proteins^42^. Notably, binding to the C-terminal peptides of the NMDA receptor subunit GluN2B^43^, which binds to the T-site, has recently been demonstrated to be extremely important for the induction of LTP^44,45^. The mechanism by which the extended structure is maintained by the CaMKIIβ subunit, as determined via HS-AFM, has the potential to contribute to prolonged exposure of the T-site, thereby promoting binding to GluN2B.

**Fig. 5:**
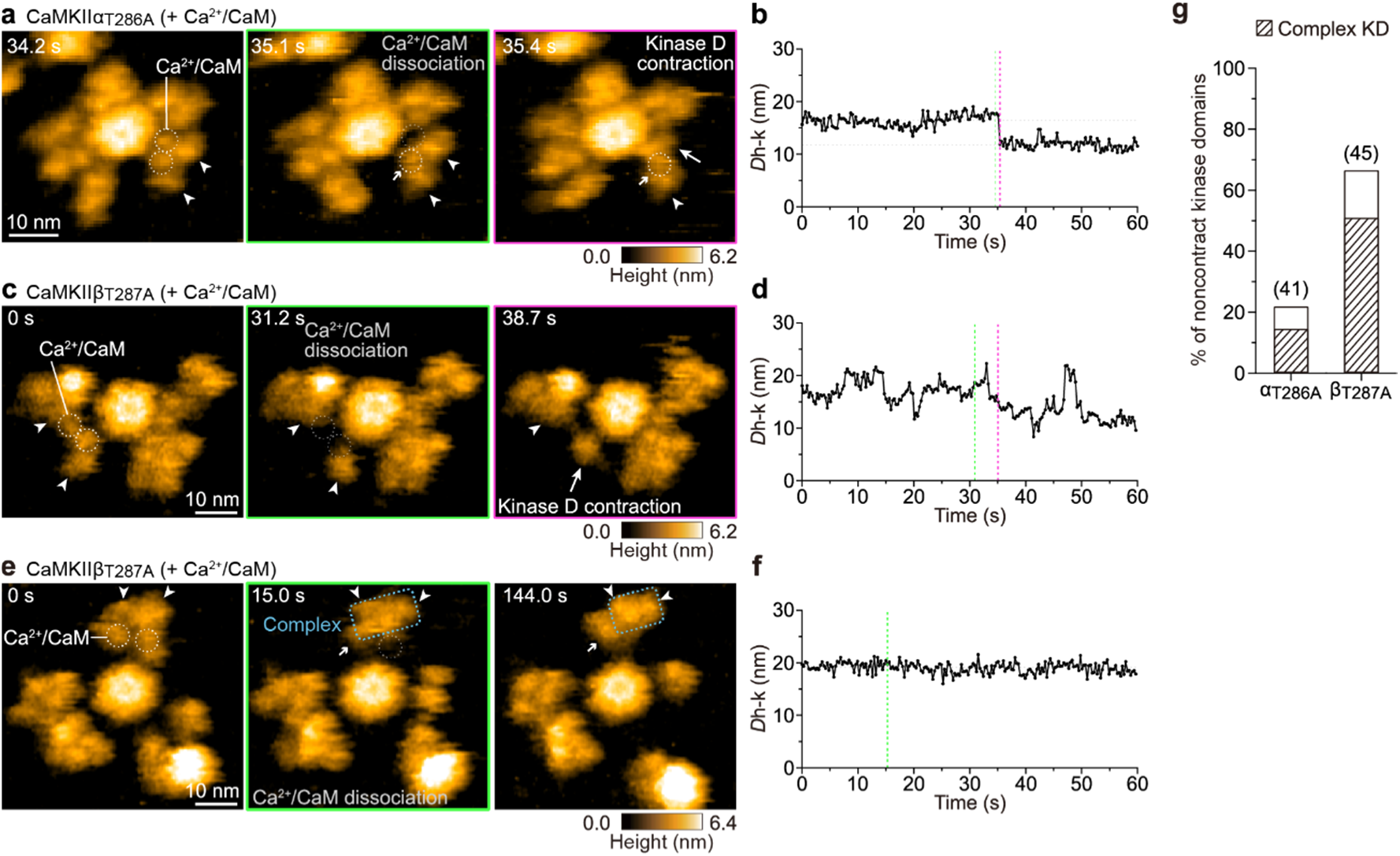
Kinase domain complexes formed in the activated state slowly degrade to the basal state after Ca^2+^/CaM dissociation. **a**,**c**,**e**, Sequential HS-AFM images of Ca²⁺/CaM-bound CaMKIIα_T286A_ (**a**) and CaMKIIβ_T287A_ (**c**, **e**) (1 mM Ca²⁺ and 800 nM CaM; see also Supplementary Video 11). **b**,**d**,**f**, Distances from the center of the hub assembly to the kinase domains (*D*_h-k_) as a function of time. The magenta and green dotted lines indicate the events of Ca²⁺/CaM dissociation and kinase domain contraction, respectively. The gray dotted lines indicate the average *D*_h-k_ before and after kinase domain contraction. **g**, Percentage of noncontracting kinase domains over 30 s after Ca²⁺/CaM dissociation. The shaded and unshaded areas represent kinase domains that formed and did not form kinase domain complexes, respectively.

## Discussion

CaMKII is a highly conserved holoenzyme that is crucial for signal transduction via Ca^2+^ signaling and has been conserved in a variety of species throughout evolution^46^. Particularly abundant in brain neurons, CaMKIIα and CaMKIIβ are present at different ratios in various brain regions^23,24^ and developmental stages^28,29^. However, structural evidence for CaMKIIα/β heterooligomers at the single-molecule level, including the localization of each subunit within the 12-meric ring structure and the functional contributions, remains unknown. In this study, we applied HS-AFM to CaMKIIβ homooligomers and CaMKII heterooligomers with an α:β ratio of 3:1, which mimics the variant ratio in the forebrain, to directly visualize phosphorylation-dependent activated states at the single-molecule level. We report the following findings. (i) Within the CaMKII 12-meric ring, variants form a mixed heterooligomer, with CaMKIIβ subunits adjacent to each other with a probability greater than 80%. (ii) In the activated state of Ca^2+^/CaM-bound CaMKII, which is followed by the phosphorylation-active state at Thr287, the kinase domain of the CaMKIIβ subunit forms a stable kinase domain complex. (iii) This kinase domain complex has low kinase activity but does not affect dephosphorylation resistance. (iv) The kinase domain complex plays a structural role in prolonging the decay time from the activated state back to the ground state.

As mentioned above, CaMKIIβ contains a binding site for F-actin in its linker region. In the inactive state, CaMKIIβ binds to F-actin, but it dissociates in the Ca^2+^/CaM-bound state^34^. This dynamic binding and dissociation of F-actin is thought to contribute to the increase in spine size during LTP^34^. In forebrain-mimicking CaMKIIα/β heterooligomers, the CaMKIIβ subunit was observed to exhibit an adjacent ring structure. To investigate the effect of this structure on F-actin binding, we performed a mathematical modeling analysis (see the Methods). Here, we used a model in which three CaMKIIβ subunits were assumed to be present within a 12-mer, with the CaMKIIβ subunits arranged either in an equally spaced configuration (subunit-dispersion model) or adjacent to each other (subunit-adjacent model) (Fig. 6a, b). The potential energy was calculated using the Lennard‒Jones potential, which is defined by the distance between the CaMKII model and the interaction point in the F-actin model as a first approximation, from which the binding probability was derived (see the Methods; Fig. 6c-f). The results indicated that the total probability of binding to F-actin in the CaMKIIβ subunit-adjacent model was approximately 22 times greater than that in the CaMKIIβ subunitdispersion model (Fig. 6g). This difference in binding probability can be attributed to the fact that the CaMKIIβ subunit-adjacent model can bind to F-actin at two points within a close distance, whereas the CaMKIIβ subunit-dispersion model can bind to F-actin at only one point (Fig. 6d, f).

**Fig. 6:**
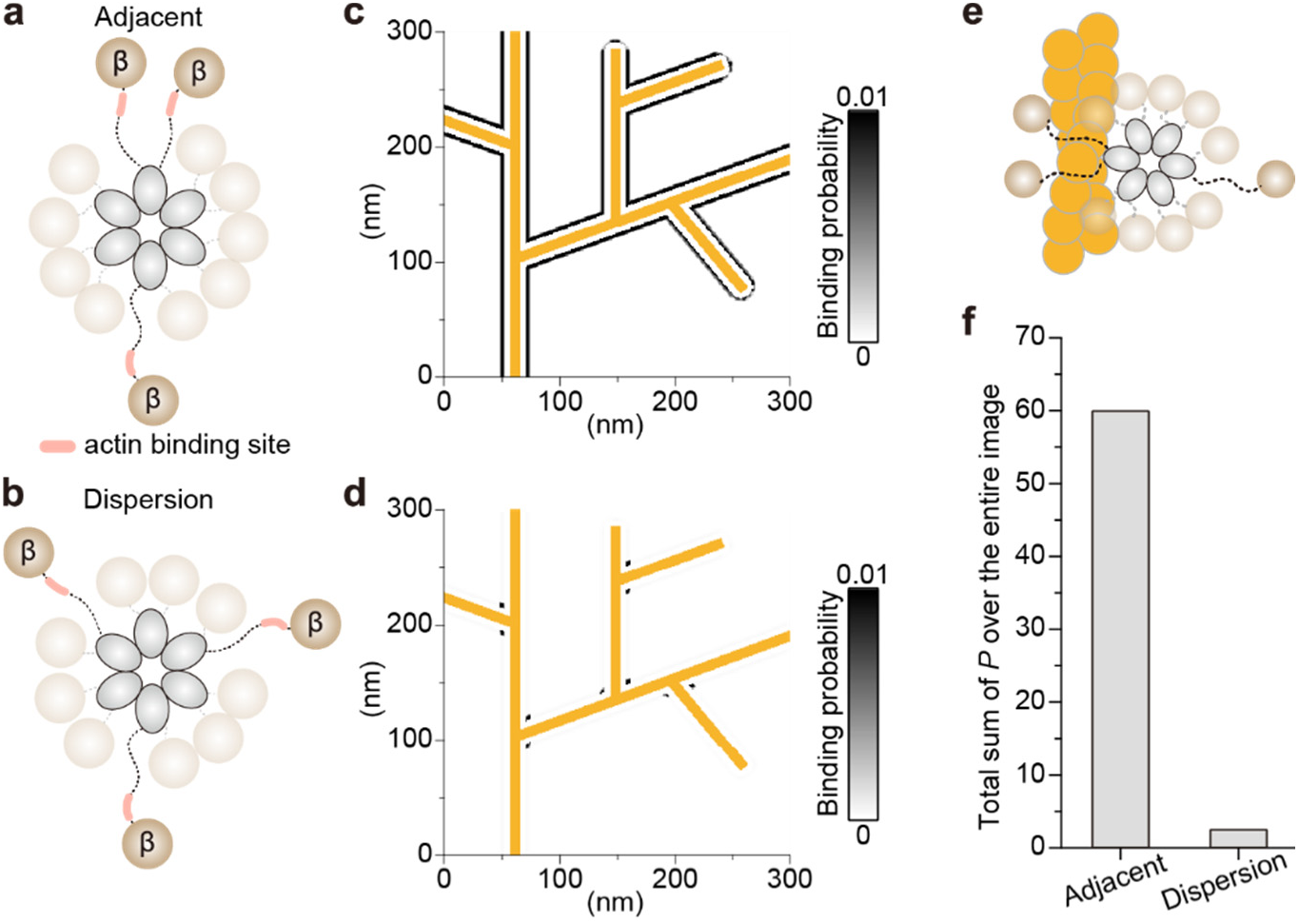
The binding of CaMKIIα/β heterooligomers to F-actin is enhanced by the adjacent ring structure of the CaMKIIβ subunits. **a**,**b** Schematic diagrams of the subunit-adjacent model (**a**) and the subunit-dispersion model (**b**). **c**,**d**, Heatmap showing the binding probability between F-actin and CaMKIIα/β heterooligomers for the subunit-adjacent model (**c**) and the subunit-dispersion model (**d**). The orange region represents the F-actin model used in this analysis, whereas the white region indicates areas where no calculations were performed. **e**, Schematic diagram of the binding pattern between F-actin and CaMKIIα/β heterooligomers in the subunit-adjacent model. **g**, Total binding probability within the analyzed region.

In the mathematical analysis of the heterooligomer model on the basis of the number of CaMKIIβ subunits, the total probability of binding in the CaMKIIβ subunit-dispersion model increased by factors of 1.2, 2.1, and 12.4 for models containing two, three, and four CaMKIIβ subunits, respectively, compared with the model with one CaMKIIβ subunit (Extended Data Fig. 10e-g). However, in the CaMKIIβ subunit-adjacent model, the total probability of binding reached 43.6-fold when there were two CaMKIIβ subunits (Extended Data Fig. 10e-g). This finding suggests that the arrangement of CaMKIIβ subunits (especially whether they are adjacent) is more critical for binding to F-actin than the total number of CaMKIIβ subunits in the 12-mer. In a previous fluorescence microscopy study that used cells, the α:β ratio of CaMKII heterooligomers that bound to F-actin with a probability of 50% was reported to be 6.5:1, indicating that for a 12-mer heterooligomer to stably bind to F-actin, it must contain at least two CaMKIIβ subunits^31^. Thus, on the basis of this mathematical model, it can be inferred that the binding strength with F-actin is greatly increased by the structure formed by adjacent CaMKIIβ subunits in the ring. Considering reports that at least two CaMKIIβ subunits are required for binding to F-actin in cells, the adjacent arrangement of CaMKIIβ subunits in the heterooligomer may play a crucial role in enabling stable binding to F-actin.

Based on these results, we propose a model for the potential activation of CaMKIIα/β heterooligomers in the spine of the forebrain (Fig. 7). In the basal state, CaMKII binds to F-actin via two adjacent CaMKIIβ subunits (Fig. 7a). Upon Ca^2+^ stimulation, Ca^2+^/CaM binds to the CaMKIIβ subunit. However, the Ca^2+^/CaM binding site competes for the F-actin binding site, resulting in dissociation of the CaMKIIα/β heterooligomer from F-actin. Concurrently, the S- and T-sites within the kinase domain become exposed. However, the CaMKIIβ subunits in the 12-mer are adjacent to one another, forming kinase domain complexes (dimers) that mask each other’s kinase active sites, leading to reduced kinase activity. Conversely, the pT287 site remains exposed, allowing contact with phosphatases, while the T-site is also accessible, facilitating binding to various proteins. Additionally, the dimeric structure of the CaMKIIβ subunits is slow to revert to the basal state after the dissociation of Ca^2+^/CaM due to the stability of the dimer, resulting in the T-site remaining exposed for an extended period. Recent reports indicate that the structural changes that lead to T-site exposure and binding to the C-terminal peptide of GluN2B are more critical for the induction of LTP than the kinase activity of CaMKII^44,45^. Collectively, these findings suggest that the prolonged active conformation induced by the adjacent CaMKIIβ heterooligomer may facilitate the binding of the C-terminal peptide of GluN2B. Thus, CaMKII forms a mixed-subunit heterooligomer in which the minimum required number of CaMKIIβ subunits (two) enables binding to F-actin. This configuration may establish a mechanism by which minimal Ca^2+^/CaM facilitates dissociation from F-actin and translocation of the remaining CaMKIIα to the vicinity of NMDAR (Fig. 7b), contributing to the diverse cellular functions of different neurons in the brain.

**Fig. 7:**
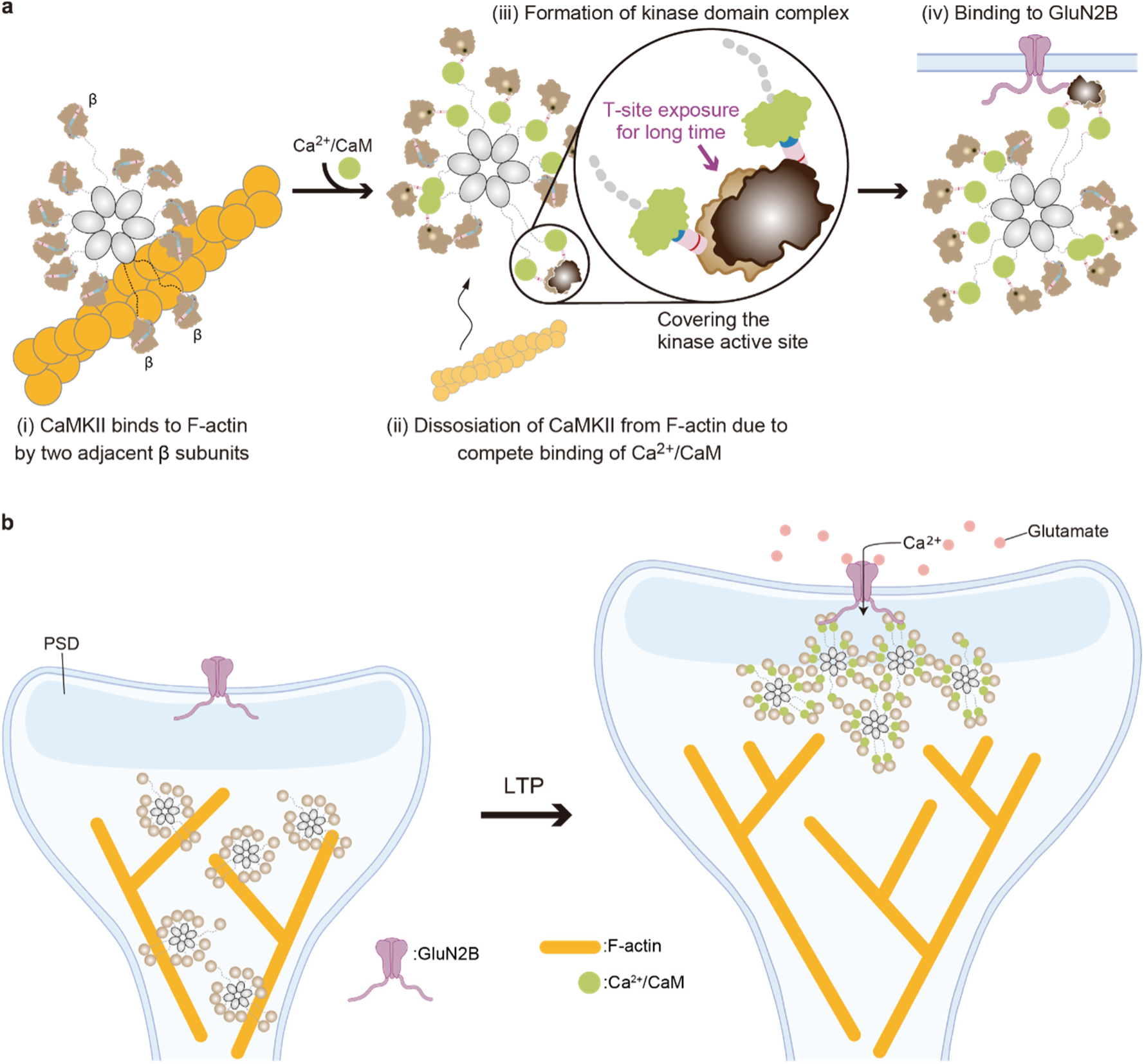
Model of the activation of CaMKIIα/β heterooligomers in the spine. **a**, Depicted the transition of binding of CaMKIIα/β heterooligomers from F-actin to GluN2B. A magnified view focusing on the kinase domain complex by CaMKIIβ subunits is shown. **b**, The inside of the spine is depicted before (left) and after (right) Ca²⁺ stimulation.

Our HS-AFM observations are limited to monitoring the dynamics of the kinase domains of CaMKIIα/β in two-dimensional space (2D) on the AFM substrate. Furthermore, direct visualization of the binding and dissociation dynamics of CaMKIIα/β heterooligomers with F-actin and GluN2B is challenging. This difficulty arises because current HS-AFM approaches are specialized for high-resolution imaging of CaMKII 12-mer ring structures. Under these conditions, CaMKII is strongly adsorbed to the AFM substrate, making visualization of its dynamics, such as protein‒protein interactions, impossible. In the near future, we will explore more refined AFM substrate conditions, such as the use of a supported lipid bilayer, to directly visualize the binding and dissociation of CaMKIIα/β heterooligomers with F-actin and GluN2B. This will contribute to our understanding of the molecular mechanisms underlying signaling processes that occur within the spine.

In conclusion, within the CaMKIIα/β heterooligomer, the α and β subunits are mixed within the multimeric ring, with β subunits found adjacent to each other. The presence of asymmetric variants within this ring facilitates binding to F-actin, and in the activated state induced by Ca^2+^/CaM binding, a stable kinase domain dimer that extends T-site exposure is formed. Therefore, direct imaging of the dynamics of the CaMKIIα/β heterooligomer via single-molecule imaging using HS-AFM shows promise for refining molecular models of CaMKII-mediated signaling mechanisms and improving our understanding of the molecular mechanisms underlying learning and memory.

## Methods

### DNA plasmids

The rat CaMKIIα, CaMKIIα_T286A_ and Calmodulin genes were prepared previously^39^. Regarding PP2A-related genes^47^, human PPP2CA, human PTPA (PPP2R4) and Mini-A (PPP2R1A) were also prepared previously^39^. Plasmids containing the CaMKIIβ gene were gifts from Tobias Meyer (Addgene plasmid #21227). sfGFP-CaMKIIβ and sfGFP-Mini-A were constructed by inserting sfGFP [ref.^48^] and the corresponding components into the modified pEGFP-C1 vector by replacing EGFP (Clontech). For all the constructs except the Calmodulin construct, 6× His/Strep tags (MDYKDDDD<u>HHHHHH</u>K<u>WSHPQFEK</u>GTGGQQMGRDLYDDDDKDLYKSGLRSRA) were fused to the N-termini of the respective genes and inserted into the modified pEGFP-C1 vector by replacing EGFP (Clontech). CaMKII mutants with point mutations were constructed using a QuikChange site-directed mutagenesis kit (Agilent Technologies, USA). For the 6× His-tagged CaM and sfGFP-fused Syntide-2 peptide (amino acid sequence PLARTLSVAGLPGKK) constructs, the respective genes were inserted into the pRSET bacterial expression vector (Invitrogen, USA). For these constructs, a 6× His tag (MRGS<u>HHHHHH</u>GMASMTGGQQMGRDLYDDDDKDRSEFG) was fused to the N-termini of the respective genes.

### Purification of Calmodulin and sfGFP-Syntide-2 from bacteria

His-tagged Calmodulin and sfGFP-fused Syntide-2 were expressed in *Escherichia coli* (DH5α) as described previously^39^. Briefly, the cell pellet was dissolved in PBS with 1% Triton X-100 and 5 mM imidazole, sonicated, and then centrifuged. The supernatant was loaded onto a Ni^2+^-nitrilotriacetate (NTA) column (HiTrap, GE Healthcare, USA). The NTA column with the bound protein was washed with 5 mM and 50 mM imidazole containing 20 mM Tris/150 mM KCl, pH 7.4 buffer, and eluted with 20 mM Tris-HCl/500 mM imidazole buffer. The concentration of the purified protein was measured using the Bradford assay (Bio-Rad, USA) with BSA as a standard. The resulting protein concentration was typically ∼500 µM in a total volume of 1000 μl. The purified proteins were stored in 20 mM Tris-HCl/150 mM KCl with 5 mM DTT at 4°C or -30°C.

### Purification of CaMKII and PP2A from HEK293 cells

All CaMKII proteins were prepared as described previously^39^. Briefly, 6× His/Strep-tagged proteins were expressed in HEK293 cells. The cells were transfected with a plasmid encoding the CaMKIIα/β genes using Lipofectamine 2000 following the manufacturer’s protocol (Thermo Fisher Scientific, USA). For the CaMKIIα/CaMKIIβ and CaMKIIα/sfGFP-CaMKIIβ complexes, two plasmids were cotransfected at a ratio of ∼3 (CaMKIIα) to 1 (CaMKIIβ or sfGFP-CaMKIIβ). One day after transfection, the cells were lysed with ice-cold lysis buffer (1% Triton X-100, 5% glycerol, 50 mM Tris, 150 mM KCl, and 4 mM EDTA, pH adjusted to 7.6) and then centrifuged at 20,000 × g for 10 min. The supernatant was loaded onto Strep-Tactin Sepharose (Nacalai, Japan), washed with 20 mM Tris-HCl/150 mM KCl, and subsequently eluted with 2.5 mM desthiobiotin/20 mM Tris-HCl/150 mM KCl, pH 7.4. The pooled proteins were further purified with an NTA column, and the protein concentration was determined using the Bradford assay. The resulting protein concentrations typically ranged from 1∼10 µM in a total volume of 500 μl. The purified proteins were stored in 20 mM Tris-HCl/150 mM KCl with 50% glycerol and 5 mM DTT, pH 7.4, at -30°C.

For PP2A, three plasmids encoding PPP2CA/PTPA/sfGFP-Mini-A genes were cotransfected at a ratio of 1:1:1. One day after transfection, the cells were lysed with ice-cold lysis buffer (1% Triton X-100, 5% glycerol, 50 mM Tris, 150 mM KCl, 0.5 mg/ml BSA, 1 mM ATP, pH adjusted to 7.6) and then centrifuged at 20,000 × g for 10 min. The supernatant was loaded onto an NTA column, and the bound protein was washed with buffer (20 mM Tris/150 mM KCl/5% glycerol/0.5 mg/ml BSA/1 mM ATP, pH 7.4) containing 50 mM imidazole and eluted with buffer containing 500 mM imidazole. The PP2A activity was determined as described previously^39^. The resulting PP2A activity was typically ∼0.2 unit/µl in a total volume of 500 μl. The purified proteins were stored in 20 mM Tris/150 mM KCl/50% glycerol/5 mM DTT/0.5 mg/ml BSA/1 mM ATP, pH 7.4, at -30°C.

### Kinase activity assay

The kinase assay was performed at 30°C, with 5 min of incubation with 50 nM concentrations of purified CaMKII proteins, 800 nM CaM, 1 mM CaCl_2_, and 1 mM ATP in reaction buffer (50 mM Tris-HCl, pH 7.4; 50 mM KCl; and 10 mM MgCl_2_). To release CaM from CaMKII, 2 mM EGTA was added, and the mixture was incubated for 5 min at 30°C and for an additional 10 min at 25°C to match the conditions used for the HS-AFM observations. For substrate phosphorylation, 1 μM sfGFP-Syntide-2 was added, and the mixture was incubated for 7 min at 25°C. For CaMKII dephosphorylation by PP2A, phosphorylated CaMKII proteins were incubated with the indicated unit of PP2A for the indicated time at 25°C. The reactions were stopped via the addition of SDS sample buffer. Western blotting was performed with the following antibodies: anti-phospho-CaMKII (Thr286) (D21E4; Cell Signaling Technology, USA), anti-pT305 (abx012403; Abbexa), anti-phospho-CaMKII (Thr306) (p1005-306; PhosphoSolutions, USA), anti-phospho-PKA substrate (RRXS*/T*) (#9624; Cell Signaling Technology), anti-His-tag (27E8; Cell Signaling Technology, USA), and HRP-anti-rabbit/mouse (Jackson Laboratory, USA).

### HS-AFM equipment

HS-AFM experiments were conducted with a custom-designed instrument in tapping mode^38,39^. The deflection of the cantilever (BL-AC10DS-A2, Olympus, Japan) was measured using an optical beam deflection sensor with an infrared (IR) laser at 780 nm and a power of 0.8 mW. The IR beam was directed onto the rear of the cantilever using a 60× objective lens (CFI S Plan Fluor ELWD 60X, Nikon, Japan). The spring constant of the cantilever was approximately 100 pN/nm, whereas its resonant frequency and quality factor in liquid were approximately 400 kHz and 2, respectively. Although the cantilever originally featured a bird beak-like triangular structure as the AFM tip, an amorphous carbon tip was synthesized on the existing AFM tip via electron beam deposition (EBD) using scanning electron microscopy (FE-SEM; Verious 5UC, Thermo Fisher Scientific, USA) to increase the spatial resolution of the HS-AFM. The length of the additional EBD tip was approximately 500 nm, and the tip apex had a radius of approximately 1 nm after further plasma etching using a plasma cleaner (Tergeo, P.I.C. Scientific, USA). The cantilever’s free oscillation amplitude was set to below 1 nm, and the setpoint amplitude for tip‒sample distance regulation was maintained at 90% of this value. Imaging acquisition and tip-scanning control were performed using laboratory-developed software written in Visual Basic .NET (Microsoft). To minimize sample damage caused by tip scanning, we employed the "only trace imaging" mode^49^. In this approach, only the trace images are acquired by scanning the tip across the sample surface, whereas no image is recorded during the retrace, in which the tip is slightly lifted away from the sample and scanned at the same speed as in the trace.

### Substrates for HS-AFM observations

Consistent with our previous study involving HS-AFM observations of CaMKIIα^39^, all the HS-AFM observations were conducted on a mica surface modified with a cationic C2 pillar[5]arene (P[5]A+) to change the surface charge from negative to positive^40,50^. Electrostatic potential mapping of CaMKIIα and CaMKIIβ revealed that the surface charge of the hub domain is partially negative, whereas the surface charge of the kinase domains is partially positive. Therefore, to prevent the inhibition of kinase domain mobility on the HS-AFM substrate, P[5]A+ was used as the HS-AFM substrate. An aqueous solution of 70 μM P[5]A+ was deposited onto a freshly cleaved mica substrate (1.0 mm in diameter; Furuuchi Chemical, Japan). P[5]A+ was incubated at room temperature on the mica surface for 15 min, and the surface was rinsed with Milli-Q water to eliminate unadsorbed P[5]A+.

### HS-AFM observations

The basal states of CaMKIIα/β heterooligomers and CaMKIIα and CaMKIIβ homooligomers, including the WT, CaMKIIα_T286A_ and CaMKIIβ_T287A_ mutants, were observed in buffer A (50 mM Tris-HCl, pH 7.4; 15 mM KCl; 10 mM MgCl_2_; and 10% glycerol) with 0.1 mM EGTA.

For the inhibitor experiment, 50 nM CaMKIIβ homooligomers were premixed at 30°C for 5 min with 50 μM bosutinib in buffer B (50 mM Tris-HCl, pH 7.4; 150 mM KCl; 10 mM MgCl_2_) supplemented with 0.1 mM EGTA. Then, HS-AFM observations were performed in buffer A with 0.1 mM EGTA and 50 μM bosutinib. Bosutinib (Cat. No. 4361) was purchased from Tocris (UK).

For Ca^2+^/CaM-bound CaMKIIα/β heterooligomers and CaMKIIβ homooligomers, 50 nM CaMKII and 800 nM CaM were premixed in buffer B with 1 mM CaCl_2_ and incubated at 30°C for 5 min. Then, HS-AFM observations were performed in buffer A with 1 mM CaCl_2_.

For Ca^2+^/CaM-bound CaMKIIα/β heterooligomers and CaMKIIβ homooligomers in the presence of ATP, we premixed 50 nM CaMKII and 800 nM CaM in buffer B with 1 mM CaCl_2_ and 1 mM ATP and incubated the mixture at 30°C for 5 min. Then, HS‒AFM observations were performed in buffer A with 1 mM CaCl_2_ and 1 mM ATP.

For CaMKIIα/β heterooligomers and CaMKIIβ homooligomers in EGTA and ATP, we premixed 50 nM CaMKII and 800 nM CaM in buffer B with 1 mM CaCl_2_ and 1 mM ATP and incubated the mixture at 30°C for 5 min. Afterward, to dissociate Ca^2+^/CaM from CaMKII, we added 2 mM EGTA and incubated the mixture at 30°C for an additional 5 min. Then, HS‒AFM observations were performed in buffer A with 2 mM EGTA and 1 mM ATP.

All HS-AFM experiments were performed at room temperature (24–26°C).

### HS-AFM image processing and data analysis

HS-AFM images were analyzed using Fiji (ImageJ) software (NIH, USA)^51^. A mean filter with a radius of 1.0 pixels was applied to each image to minimize noise. To correct drift in sequential images, the Template Matching and Slice Alignment plugin for ImageJ was utilized. The coordinates of the hub assembly center were determined using the Trainable Weka Segmentation plugin^52^. The MTrackJ plugin^53^ for ImageJ was used for manual tracking of the coordinates corresponding to the highest pixel for each kinase domain across all CaMKII protein constructs under various HS-AFM experimental conditions.

The motion of each kinase domain for the CaMKIIα/β heterooligomer (56,856 points in 4,738 frames of 528 kinase domains in 44 oligomers), the CaMKIIβ homooligomer (18,000 points in 1,500 frames of 120 kinase domains in 10 oligomers), CaMKIIβ + bosutinib (24,024 points in 2,002 frames of 168 kinase domains in 14 oligomers), CaMKIIβ + ADP (19,608 points in 1,634 frames of 132 kinase domains in 11 oligomers), CaMKIIβ + ATP (17,531 points in 1,473 frames of 119 kinase domains in 10 oligomers), and CaMKIIβ in EGTA/ATP after Ca^2+^/CaM/ATP stimulation (25,512 points in 2,126 frames of 192 kinase domains in 16 oligomers) was analyzed. The motion of the Ca^2+^/CaM-binding kinase domains in CaMKIIβ with ATP (18,000 points in 1,500 frames of 120 kinase domains in 10 oligomers) and without ATP (18,000 points in 1,500 frames of 120 kinase domains in 10 oligomers) was analyzed.

The motion of each kinase domain of CaMKIIα_T286A_ (7,767 points of 41 kinase domains in 22 oligomers) and CaMKIIβ_T287A_ (8,393 points of 45 kinase domains in 32 oligomers) was analyzed.

HS-AFM experiments were independently repeated at least three times with similar results.

### Analysis of kinase domain trajectories

To quantify the trajectory of a single kinase domain, the distance from the center of the hub assembly to the kinase domain (*D*_h-k_) was computed as follows:

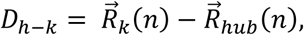

Where 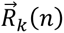 is the position of the kinase domain in video frame (*n*) and where 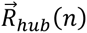 is the position of the center of the hub assembly in video frame (*n*).

To characterize a single kinase domain, the radius of gyration, 𝑅_g_, of the kinase domain trajectory was computed as the root-mean-square displacement from its average position:

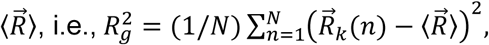

where 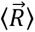 is the average position of a single kinase domain. *R_g_* was calculated from HS-AFM videos over a duration of 45 s, with a temporal resolution of 0.3 s per frame.

### Distinguishing CaMKIIα and CaMKIIβ subunits in CaMKIIα/β heterooligomers

First, the distribution of *D*_h-k_ for the CaMKIIα and CaMKIIβ homooligomers indicated that kinase domains with *D*_h-k_ values greater than 14 nm could be identified as CaMKIIβ subunits. Consequently, 97.9% (*n* = 144 kinase domains) of the kinase domains in CaMKIIα homooligomers were identified as CaMKIIα subunits, whereas 66.7% (*n* = 120 kinase domains) of the kinase domains in the CaMKIIβ homooligomers were classified as CaMKIIβ subunits. This suggests that 33.3% of CaMKIIβ subunits would be misidentified as CaMKIIα subunits if assessed solely on the basis of *D*_h-k_ values. To account for the remaining CaMKIIβ subunits, we introduced an additional criterion: kinase domains with an *R*_g_ value greater than 2.8 nm were classified as CaMKIIβ subunits, even if their *D*_h-k_ value was less than 14 nm. This *R*_g_ value threshold of 2.8 nm was estimated on the basis of the distribution of *R*_g_ values for CaMKIIα and CaMKIIβ homooligomers. Using this *R*_g_ criterion, 91.7% of the kinase domains in the CaMKIIα homooligomers were identified as CaMKIIα subunits, whereas 61.7% of the kinase domains in the CaMKIIβ homooligomers were identified as CaMKIIβ subunits. By combining these two criteria (first, *D*_h-k_ values over 14 nm, and second, *R*_g_ values greater than 2.8 nm, even if the *D*_h-k_ value was below 14 nm), 98.6% of the kinase domains in the CaMKIIα homooligomers were identified as CaMKIIα subunits, whereas 80.8% of the kinase domains in the CaMKIIβ homooligomers were classified as CaMKIIβ subunits. Notably, an error of approximately 19.2% (2.3 subunits out of a 12-mer) could result in the misidentification of CaMKIIβ subunits as CaMKIIα subunits, suggesting that the actual number of CaMKIIβ subunits in CaMKIIα/β heterooligomers is likely somewhat greater. We analyzed 44 CaMKIIα/β heterooligomers. Among these, two oligomers contained no CaMKIIβ subunits, whereas five oligomers contained one CaMKIIβ subunit each. Of the remaining 37 CaMKIIα/β heterooligomers, 31 oligomers exhibited adjacent CaMKIIβ subunits. This finding indicates that CaMKIIβ subunits are adjacent to one another, with a probability of 83%, in CaMKIIα/β heterooligomers within the 12-meric ring.

### Analysis of kinase domain complexes

Kinase domain complexes were defined on the basis of the distance between kinase domains and their mobility. First, the distance between the maximum heights of each kinase domain (*D*_k-k_ in Fig. 3f) was measured and found to be shorter than the length of the major axis of the kinase domains. The length of the major axis of the kinase domains was calculated in regions greater than 2 nm in height from the AFM substrate and recorded as follows: 6.0 ± 0.7 nm (120 kinase domains in 10 oligomers) in the basal state, 6.4 ± 0.9 nm (120 kinase domains in 10 oligomers) in the Ca^2+^/CaM binding state, 6.2 ± 0.8 nm (120 kinase domains in 10 oligomers) in the Ca^2+^/CaM+pT287 state, and 6.6 ± 1.0 nm (192 kinase domains in 16 oligomers) in the pT287/pT306/pT307 state. Next, to exclude kinase domain complexes consisting of two kinase domains located close to each other in each HS-AFM frame, we used the mobility of each kinase domain to define the kinase domain complex. For example, if two kinase domains were positioned adjacent to each other and the distance between them exceeded the length of the major axis of the kinase domains, yet the kinase domains moved together in the same direction, we classified these kinase domains as being in a complex state. Conversely, if the distance between two kinase domains was less than or equal to the length of the major axis of the kinase domains but they did not move in the same direction, these kinase domains were not classified as being in a complex state.

For the dwell time analysis of the kinase domain complex, only cases in which the complex was maintained for more than 10 frames (0.3 [s] × 10 [frames] = 3 [s]) were counted. Additionally, when the distance between the kinase domains exceeded the length of the major axis of the kinase domain for 5 frames, it indicated the release of the kinase domain complex. A total of 10 oligomers (1,500 frames) in the basal state, 10 oligomers (1,500 frames) in the Ca^2+^/CaM-bound state, 10 oligomers (1,500 frames) in the Ca^2+^/CaM + pT287 state, and 9 oligomers (1,350 frames) in the pT287/pT306/pT307 state were analyzed to evaluate the dwell time of the kinase domain complex.

### Contraction analysis of the kinase domains of CaMKIIα_T286A_ and CaMKIIβ_T287A_

We analyzed the kinase domains from which Ca^2+^/CaM dissociated during HS-AFM scanning. The *D*_h-k_ values of these kinase domains were measured for approximately 100 frames before and after the dissociation of Ca^2+^/CaM. The dissociation of Ca^2+^/CaM was detected manually through frame-by-frame comparisons of the HS-AFM images. We defined contraction of the kinase domains to the autoinhibitory state as a decrease in the *D*_h-k_ value of approximately 2.0 nm or more. Kinase domains that did not exhibit any decrease in *D*_h-k_ values within 30 s after Ca²⁺/CaM dissociation were classified as "noncontracted kinase domains".

### Estimation of the binding of CaMKIIα/β heterooligomers to F-actin

A model structure of F-actin was generated in a 320 × 320 nm² area. F-actin was represented on the basis of its presence or absence in 1 × 1 nm² regions. The model structure of F-actin was constructed on the basis of observations from previous studies^54^.

The diameter of F-actin was set to 9 nm and the branching angle was set to 70° (ref.^56^). The generated structure is shown in Extended Data Figure 10. The position of the interaction site in the CaMKIIβ subunits was set at the midpoint of the linker (Extended Data Fig. 10a)^35,57^. The coordinates of the linker’s midpoint were geometrically calculated using the diameter of the hub domain (11 nm)^21^, the diameter of the kinase domain (4.5 nm) and the hub-to-kinase center distance (15.2 nm, obtained from HS-AFM observations; . We confirmed the binding mode of the CaMKIIβ homooligomer to F-actin by HS-AFM (Extended Data Fig. 10b).

The potential energy between the CaMKIIβ subunits and F-actin was calculated using the shifted Lennard‒Jones potential as a first approximation (Extended Data Fig. 10c,d)^58^. This potential was selected to ensure that the potential energy was minimized when the CaMKIIβ subunits came into contact with F-actin. The potential energy (*E*) was calculated using the following equation:

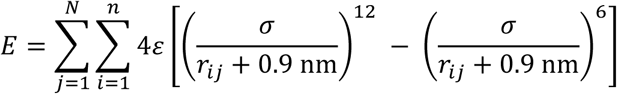

where *r_ij_* represents the distance between the interaction sites on CaMKIIβ subunits and each block of F-actin in the xy-plane. *n* denotes the number of interaction sites in the CaMKIIα/β heterooligomer, and *N* represents the number of grid points where F-actin is present. If there were multiple interaction sites on the CaMKIIα/β heterooligomer (e.g., when two or more CaMKIIβ subunits are present), the total potential energy was calculated by summing the potential energies of all interaction sites with F-actin. The parameter *σ* was set to 1 nm to ensure a wider potential well. Owing to the denominator of the equation (*r_ij_* + 0.9 nm), the potential energy becomes zero when *r_ij_* = 0.1 nm. Thus, the energy is zero when the interaction sites on CaMKIIβ are in direct contact with F-actin. The locations where CaMKIIα/β heterooligomers can bind were considered to be within the range of 5 nm < *R* ≤ 11 nm, where *R* is the distance between the center of the CaMKIIα/β heterooligomers and the nearest block of F-actin. Considering the diameter of the hub domain of the CaMKIIα/β heterooligomers (11 nm), the lower limit (*R* > 5 nm) was set to prevent the overlap of the hub domain with F-actin, as observed in the HS-AFM images (Extended Data Fig. 10b). The upper limit (*R* ≤ 11 nm) was chosen because beyond this distance, the absolute value of the potential energy becomes very small and is negligible, which was also helpful in reducing the computational time.

The *ε* used in the potential energy calculation was determined to reproduce the binding constant *K_a_* = 1.0×10⁵ M⁻¹ reported in previous studies^59^. The binding constant *K_a_* in this estimation was calculated using the following equations :

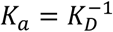

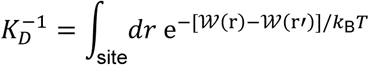

where *W(r)* represents the potential energy at position *r* and where *r′* is the reference position in the bulk. In this estimation, *W*(*r*′) is naturally zero. *k*_B_ and *T* represent the Boltzmann constant and temperature, respectively. The temperature was set to 298 K. The integration range was defined as a 6 × 11 nm^2^ region, where at least one CaMKIIα/β heterooligomer is expected to be adjacent to F-actin. This range was determined on the basis of the diameter of the hub domain (11 nm): 6 (= 11 − 5) nm vertically and 11 nm horizontally from F-actin, in which 5 nm is used to exclude the overlapping regions. The *ε* that reproduced the experimental *K_a_* value was 1.73 kcal/mol.

To obtain the optimal binding angle of the CaMKIIα/β heterooligomer, the potential energy was calculated by rotating the CaMKIIα/β heterooligomer up to 355 degrees at 5-degree intervals to find the conformation with the lowest energy. A heatmap was generated using the potential energy at the optimal angle. Margins of 10 nm were removed from all sides to exclude cases where the interaction sites of CaMKIIβ subunits were outside the calculation area. As a result, the final heatmap covered a 300 × 300 nm² region.

To calculate the binding probability (*P_a_*) from the potential energy, the following equation was used^62^:

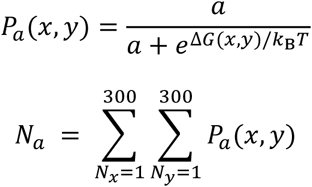

where *N_a_* is the number of bound CaMKIIα/β heterooligomers and where *N_x_* and *N_y_* are the numbers of blocks in the x and y directions in the model, respectively. Δ*G*(*x*, *y*) was considered to be the potential energy at the position. A parameter *a* was determined so that the estimated line density of bound CaMKII, *ρ* = *N_a_*/*l*, where *I* is the perimeter length of F-actin, reproduces the experimental line density (0.0278 nm^−1^)^33^. The obtained value of *a* was 2.1088×10^-6^.

### Quantification and statistical analysis

All statistical analyses were performed using Igor Pro 9 software (WaveMetrics, USA) or OriginPro2025 (OriginLab, USA). A significance level of α = 0.05 was applied for all analyses, and *P* values were adjusted for multiple comparisons where relevant. Unless otherwise stated, the normality of the data distribution was tested by the Shapiro‒Wilk test. If the normality assumption failed, the Kruskal‒Wallis test and Mann‒Whitney *U* test were used for comparisons between two groups. For comparisons involving multiple conditions, one-way ANOVA and the Kruskal‒Wallis test were used to analyze multiple groups with a single independent variable. As a follow-up test to the Kruskal‒Wallis test, Dunn’s test was used to compare every mean with every other mean. The Bonferroni correction was used as the follow-up test for one-way ANOVA. The tests used are indicated in the figure captions. *P* > 0.05, not significant (NS); \**P* < 0.05, \*\**P* < 0.01, and \*\*\**P* < 0.001 were considered to indicate statistical significance. The data are presented as the means ± SDs.

### Use of large language models

ChatGPT was used to proofread the text.

## Acknowledgments

We thank R. Yasuda (Max Planck Florida Institute) and T. Ando (Kanazawa University) for providing the HS-AFM-related apparatus; Y. Mikami and Y. Okada (Kanazawa University) for collecting and analyzing the HS-AFM data. This research was supported by the World Premier International Research Center Initiative (WPI), MEXT, Japan (to M.S.), JSPS KAKENHI grant numbers JP24K21942 (to M.S.) and JP25H00972 (to M.S.), JP22H04926 Advanced Bioimaging Support (ABiS) (to M.S.), 23H0424 (to H.M.), 24H01298 (to H.M.), the Mochida Memorial Foundation for Medical and Pharmaceutical Research (to M.S.), the Uehara Memorial Foundation (to M.S.), the Naito Foundation (to M.S.), and JST/CREST (JPMJCR1762 to N.K. and H.F.).

## Author contributions

Conceptualization: M.S.; methodology: K.M., T.S., K.U., N.K., and M.S.; software: K.M., H.F., and M.S.; validation: K.M., T.S., M.I., Y.N., A.S., H.M., and M.S.; formal analysis: K.M., T.S., A.S., H.F. and M.S. ; investigation: K.M., T.S., M.I., Y.N., A.S., H.M., and M.S.; resources: T.K., T.O., H.M., and M.S.; data curation: H.M., and M.S.; writing – original draft: M.S.; writing – review & editing: all authors; visualization: K.M., H.M., and M.S.; supervision: T.S., A.S., H.M., and M.S.; project administration: M.S.; funding acquisition: H.M., and M.S.

## Ethics declarations

The authors declare that they have no competing interests.

---

**Extended Data Fig. 1:**
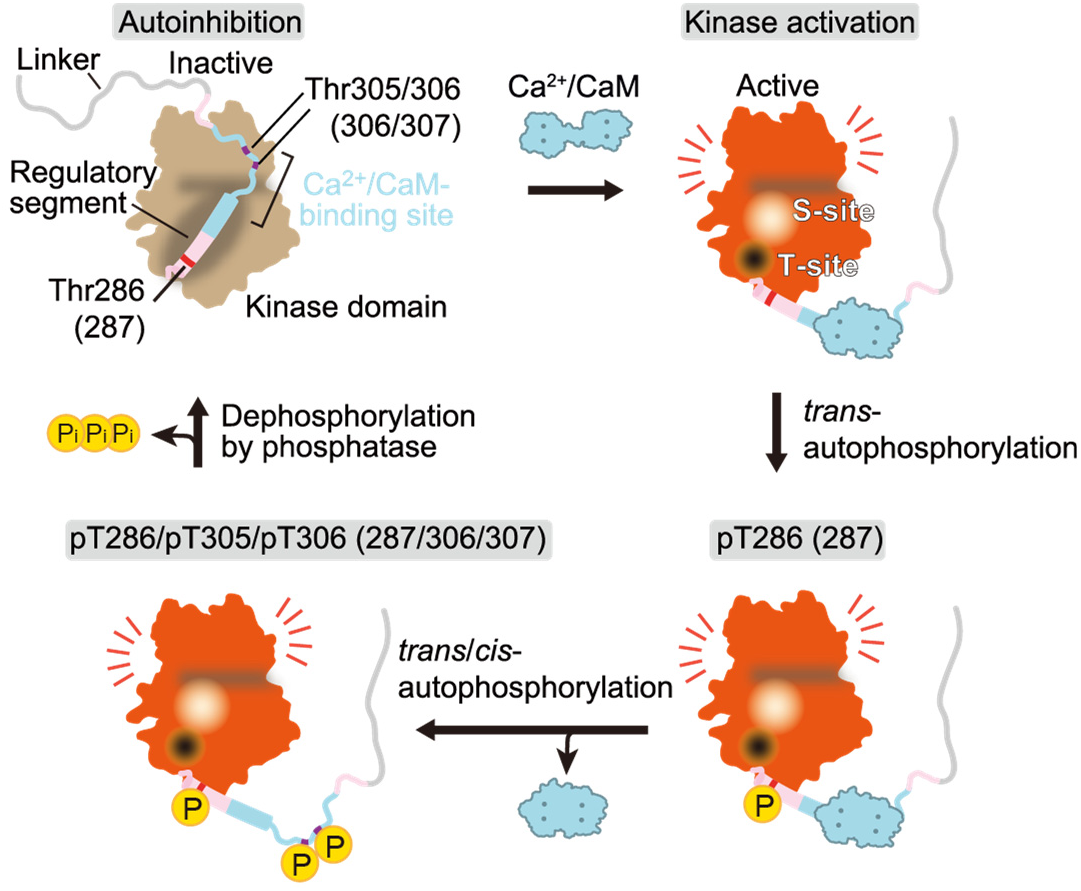
Activation state of a kinase domain in CaMKII. **a**, Illustration of the state change of CaMKII, focusing on a single kinase domain. In the inactive state, the kinase domain is autoinhibited by the regulatory segment (autoinhibition). The binding of Ca^2+^/CaM leads to the release of this regulatory segment, revealing the substrate-binding site (S-site; orange) and the GluN2B-binding site (T-site; black) (i.e., kinase activation). The concomitant activation of adjacent kinases results in autophosphorylation at Thr286 (Thr287) (autophosphorylation at pT286 (pT287)). Following the dissociation of Ca^2+^/CaM, autophosphorylation occurs at Thr305/306 (Thr306/307). Ultimately, a protein phosphatase dephosphorylates CaMKII, returning it to the inactive state. The numbers in parentheses refer to CaMKIIβ.

**Extended Data Fig. 2:**
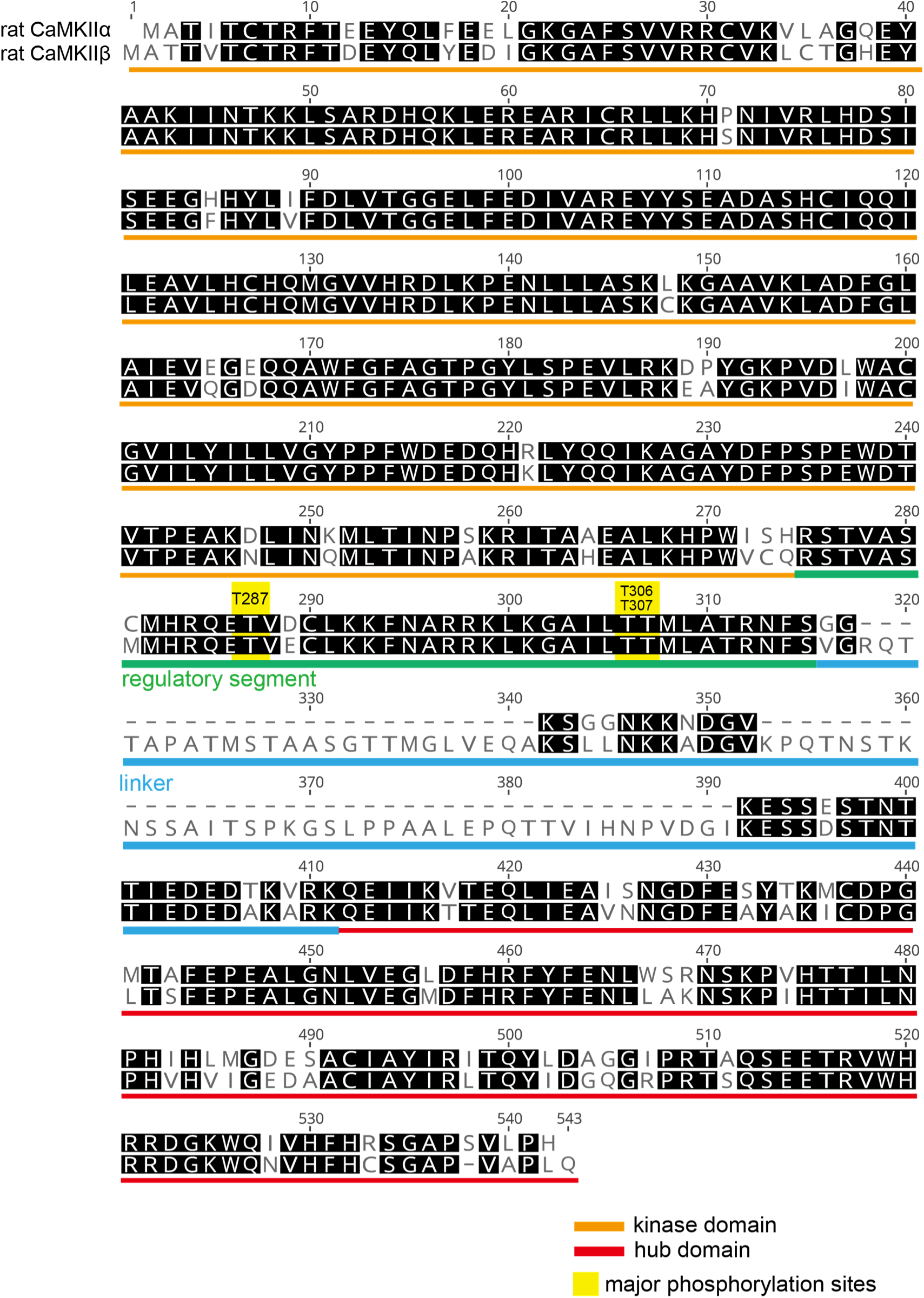
Sequence alignment of the rat CaMKIIα and CaMKIIβ used in this study. rat CaMKIIα (*Rattus norvegicus*, Accession#: NP_037052.1) and rat CaMKIIβ (*Rattus norvegicus*, Accession#: NP_ 001035813.1).

**Extended Data Fig. 3:**
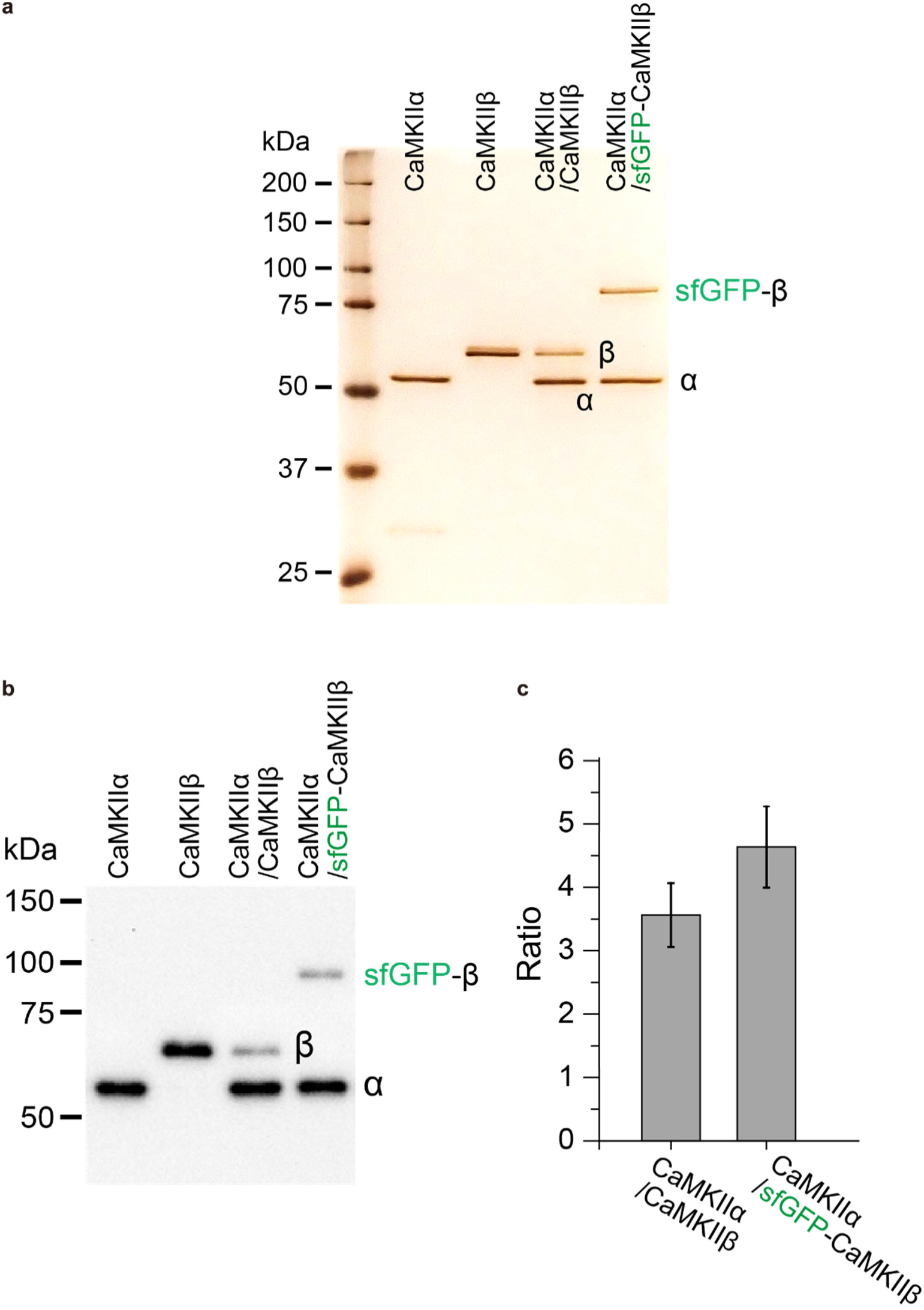
Biochemical validation of purified CaMKII proteins. **a**, The purity of CaMKII proteins was confirmed by silver staining. CaMKII holoenzymes in the absence of ADP/ATP were purified from HEK-293 cells via two-step purification with His and Strep tags. **b**,**c**, The ratio of CaMKIIα to CaMKIIβ in the purified proteins containing both CaMKIIα and CaMKIIβ was determined by western blotting (**b**), and the band intensity was quantified (**c**).

**Extended Data Fig. 4:**
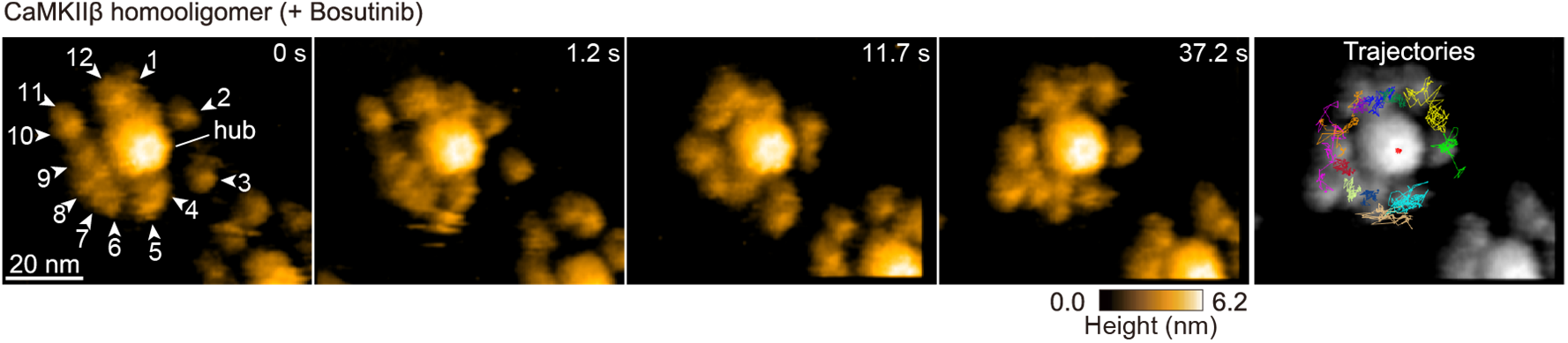
The addition of bosutinib reduces the mobility of the kinase domain in CaMKIIβ. Sequential HS-AFM images of CaMKIIβ in the presence of 50 μM bosutinib (see also Supplementary Video 2). The white arrowheads indicate kinase domains, each arbitrarily numbered. Frame rate, 3.3 frames/s. The trajectories of the kinase domains and the center of the hub assembly (red in the center) were tracked for approximately 40 s.

**Extended Data Fig. 5:**
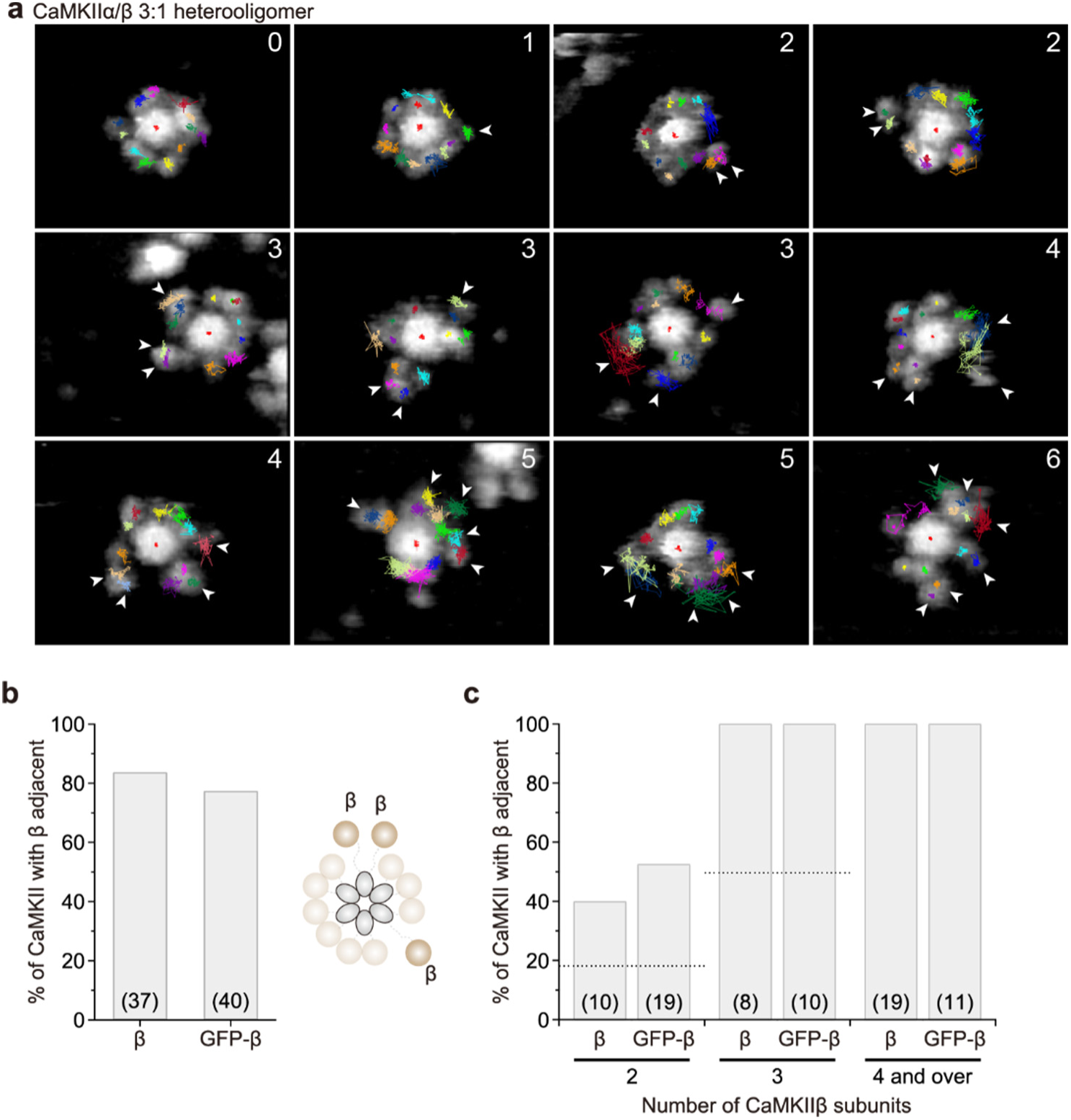
The CaMKIIα/β heterooligomer comprises a mixture of CaMKIIα and CaMKIIβ subunits in the 12-meric ring, with the CaMKIIβ subunits positioned adjacent to each other. **a**, Trajectories of the kinase domains and the center of the hub assembly (red in the center) of the CaMKIIα/β heterooligomer (approximately 30 s). Frame rate, 3.3 frames/s. The white arrowheads indicate kinase domains derived from CaMKIIβ subunits. The numbers in the upper right indicate the number of CaMKIIβ subunits (see the Methods for the criteria used to determine the number of CaMKIIβ subunits). **b**, Percentages of neighboring CaMKIIβ and GFP-CaMKIIβ subunits within a CaMKIIα/β 12-meric ring. The numbers in parentheses indicate the number of analyzed oligomers (also in **c**). **c**, Percentages of neighboring CaMKIIβ subunits and GFP-CaMKIIβ subunits within a CaMKIIα/β 12-meric ring according to the number of CaMKIIβ subunits. For two or three subunits, probabilities calculated as random chance are shown as dotted lines.

**Extended Data Fig. 6:**
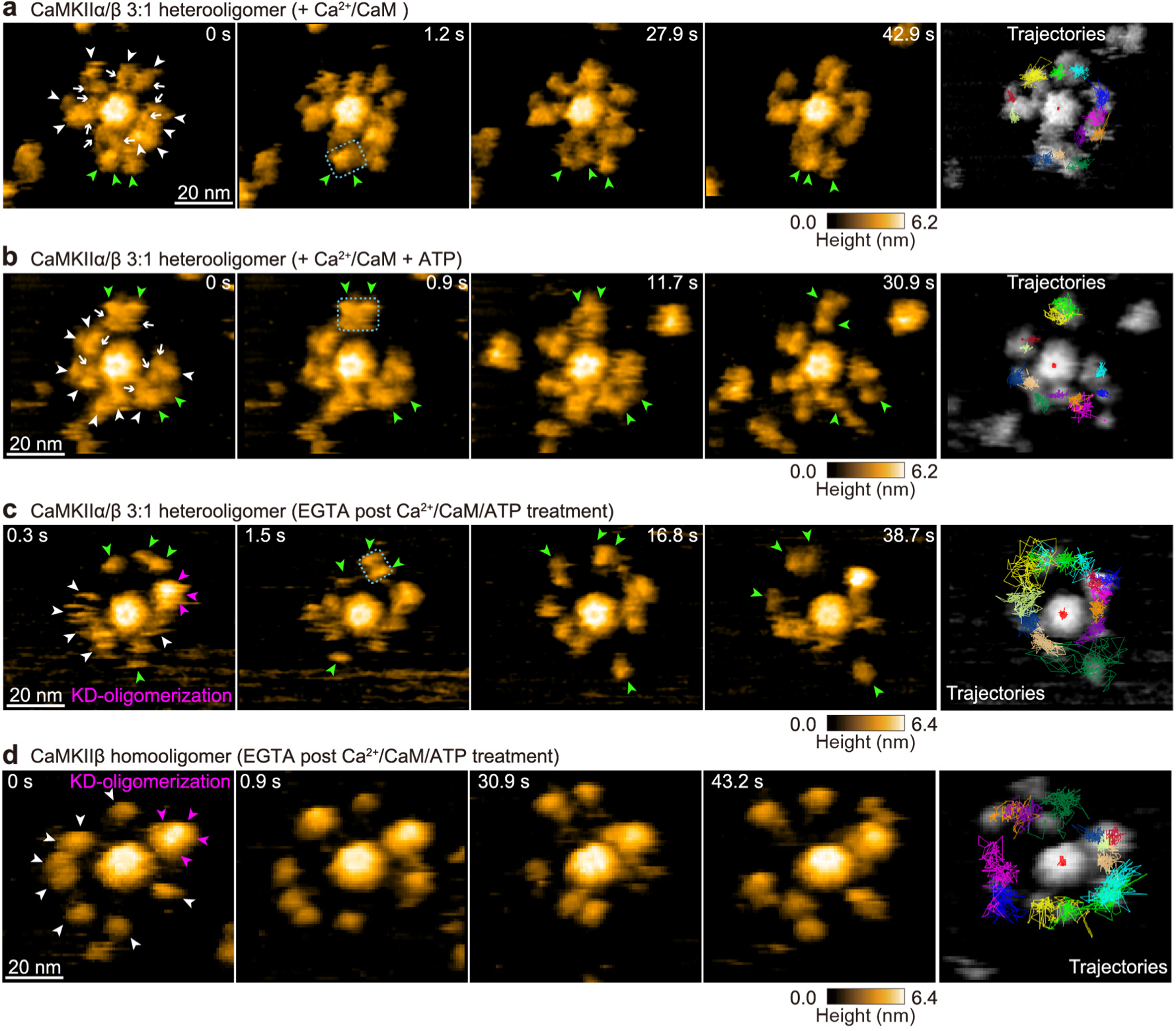
HS-AFM observations of the activation states of the CaMKIIα/β heterooligomer. **a-d**, Sequential HS-AFM images of Ca²⁺/CaM-bound CaMKIIα/β heterooligomers (**a**; 1 mM Ca^2+^, 800 nM CaM; see also Supplementary Video 6), pT286(287) CaMKIIα/β heterooligomers (**b**; 1 mM Ca^2+^, 800 nM CaM, and 1 mM ATP; see also Supplementary Video 8), pT286(287)/pT305(306)/pT306(307) CaMKIIα/β heterooligomers (**c**; see also Supplementary Video 10), and pT287/pT306/pT307 CaMKIIβ (**d**; see also Supplementary Video 9). The white arrows indicate Ca^2+^/CaM bound to the kinase domains (see the Materials for the criteria used to determine the CaMKIIβ subunits). The white and green arrowheads indicate kinase domains derived from the CaMKIIα and CaMKIIβ subunits, respectively. The blue dotted squares indicate a kinase domain complex. Magenta arrowheads indicate KD-oligomerization. Frame rate, 3.3 frames/s. The trajectories of the kinase domains and the center of the hub assembly (red in the center) were tracked for approximately 45 s.

**Extended Data Fig. 7:**
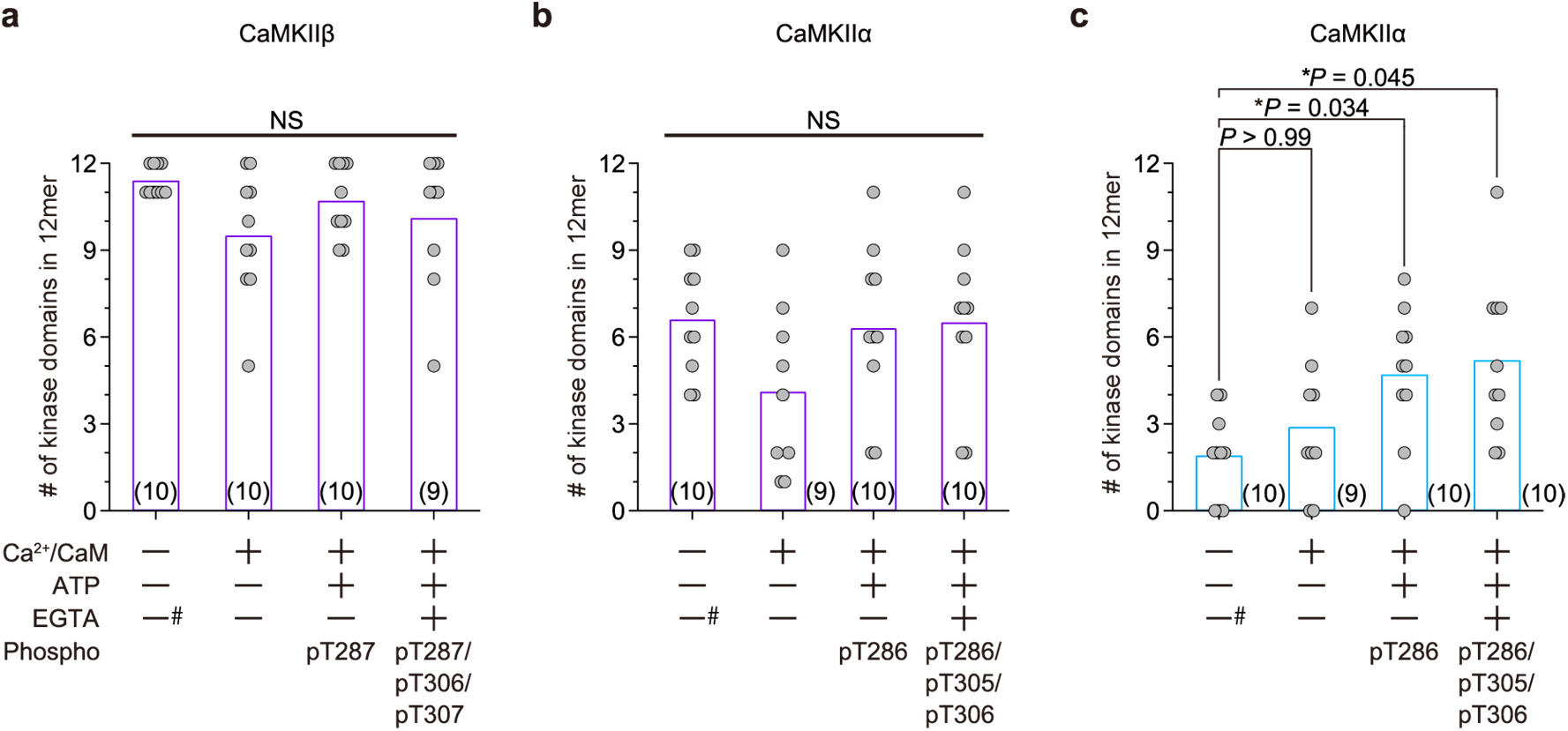
The kinase domains of CaMKIIβ form a kinase domain complex in the 12-meric ring. **a**,**b**, The number of kinase domains that formed complex states in the 12-meric ring per CaMKIIβ (**a**) and CaMKIIα (**b**) homooligomer under the indicated conditions. The number in parentheses is the number of holoenzymes used in the analysis (Kruskal‒Wallis test). **c**, The number of kinase domains that remained in the complex state for more than 45 s per 12-meric CaMKIIα homooligomer under the indicated conditions. The number in parentheses is the number of holoenzymes used in the analysis (Kruskal‒Wallis test with Dunn’s post hoc test). #: 0.1 mM EGTA

**Extended Data Fig. 8:**
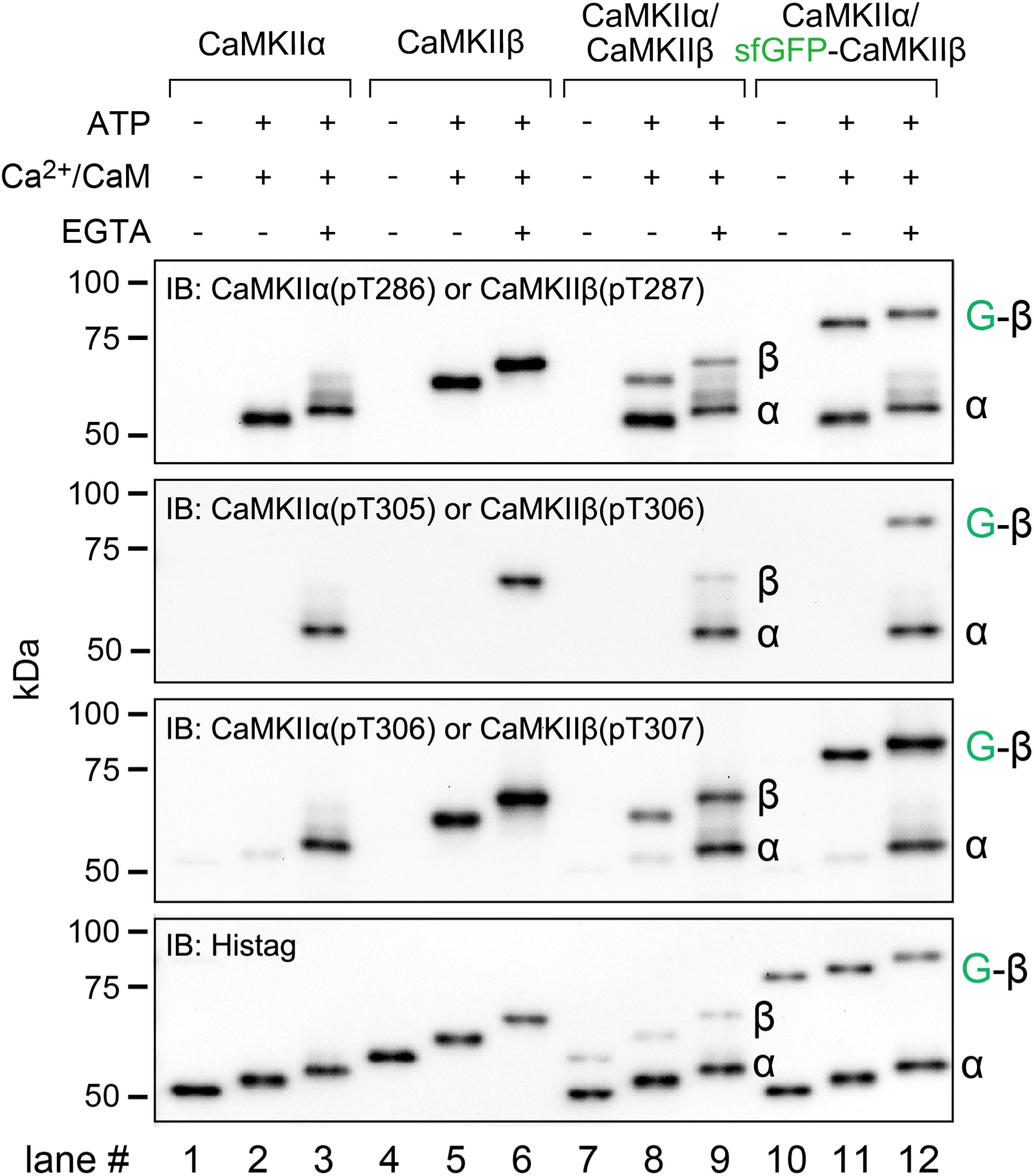
Phosphorylation of purified CaMKII. (lanes #1, 4, 7, 10) Purified proteins were loaded without activation. (lanes #2, 5, 8, 11) CaMKII proteins (50 nM) were incubated with 800 nM CaM, 1 mM CaCl_2_, and 1 mM ATP in reaction buffer (50 mM Tris-HCl, pH 7.4; 50 mM KCl; and 10 mM MgCl_2_) at 30°C for 5 min. This protocol selectively induces phosphorylation at Thr286 of CaMKIIα and Thr287/Thr307 of CaMKIIβ. (lanes #3, 6, 9, 12) Ca^2+^/CaM was dissociated by incubation with 2 mM EGTA for 5 min at 30°C and an additional 10 min at 25°C. This protocol induces the phosphorylation of CaMKIIα at Thr286/305/306 and CaMKIIβ at Thr287/306/307. The amount of protein loaded was assessed with an anti-His tag antibody.

**Extended Data Fig. 9:**
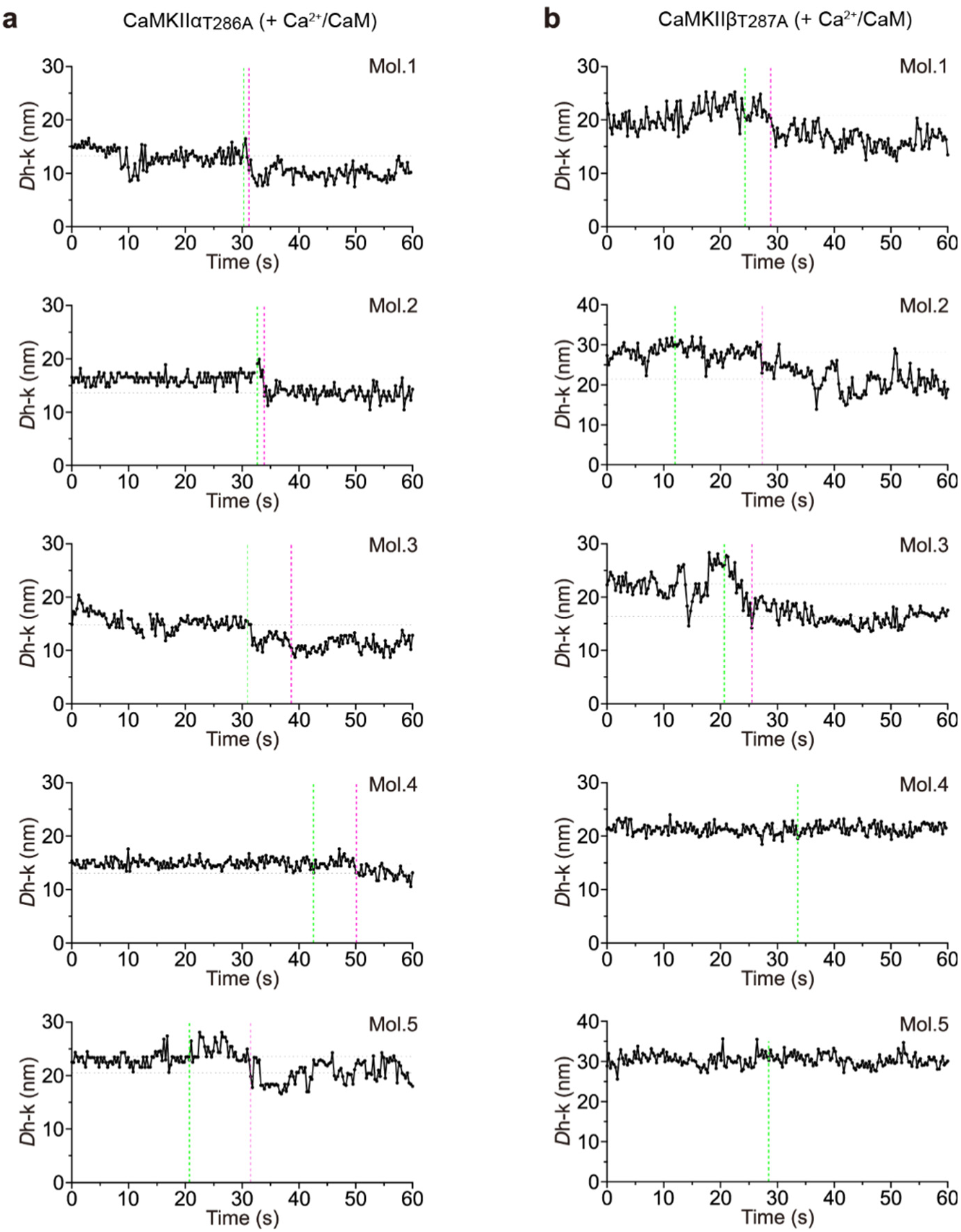
Kinase domain complexes formed in the activated state exhibit slow structural decay to the basal state after Ca^2+^/CaM dissociation. **a**,**b**, Five representative distances from the center of the hub assembly to kinase domains (*D*_h-k_) as a function of time for CaMKIIα_T286A_ (**a**) and CaMKIIβ_T287A_ (**b**). The magenta and green dotted lines indicate the events of Ca²⁺/CaM dissociation and kinase domain contraction, respectively. The gray dotted lines indicate the average *D*_h-k_ before and after kinase domain contraction.

**Extended Data Fig. 10:**
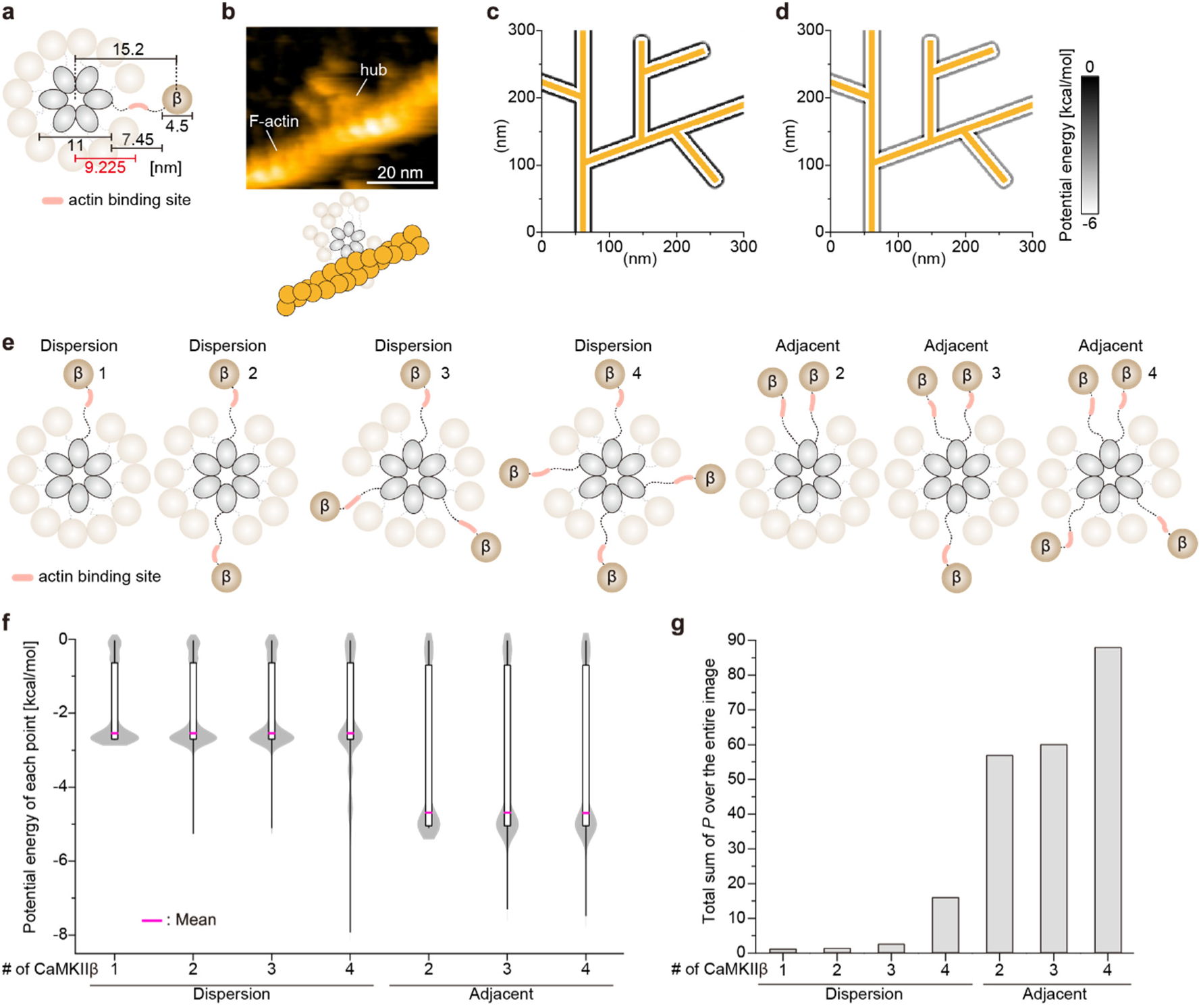
The binding strength of CaMKIIα/β heterooligomers to F-actin is enhanced by the adjacent ring structure of the CaMKIIβ subunits. **a**, Position of the interaction point with F-actin on the CaMKIIβ linker. The interaction point is set at the midpoint of the linker (pink area). **b**, HS-AFM image of a complex of the CaMKIIβ homooligomer and F-actin. **c**,**d** Heatmap of the potential energy when the kinase domains of CaMKIIβ subunits are adjacent (**c**) and evenly dispersed (**d**). The F-actin model is shown in orange. **e**, Models of CaMKIIα/β heterooligomers with different numbers of CaMKIIβ subunits. The angles between CaMKIIβ subunits are 180°, 60° and 90° for the subunit dispersion model and 30°, 30° and 165°, 30° and 120° and 90° for the subunit-adjacent model. **f**, Potential energy at each pixel for different CaMKIIα/β heterooligomer models. **g**, Total binding probability of the CaMKII heterooligomer and F-actin in the different CaMKIIα/β heterooligomer models.

