## Supplemental video legends for "Structural dynamics of mixed-subunit CaMKIIα/β heterododecamers filmed by high-speed AFM"

**Supplementary Video legends**

**Supplementary Video 1. HS-AFM videos of three representative CaMKIIβ homooligomers on a P[5]A+-modified mica surface.** The trajectories of the kinase domains are shown on the right. Image size, 360 × 98 pixels$^2$ (acquired in HS-AFM observations); scan area, 80 × 68 or 69 nm$^2$; frame rate, 3.3 fps.

**Supplementary Video 2. HS-AFM videos of three representative CaMKIIβ homooligomers with 50 μM bosutinib on a P[5]A+-modified mica surface.** The trajectories of the kinase domains are shown on the right. Image size, 240 × 96 pixels$^2$ (acquired in HS-AFM observations); scan area, 80 × 64 nm$^2$; frame rate, 3.3 fps.

**Supplementary Video 3. HS-AFM videos of three representative CaMKIIα/β 3:1 heterooligomers on a P[5]A+-modified mica surface.** The trajectories of the kinase domains are shown on the right. Green arrowheads indicate the kinase domains of CaMKIIβ subunits. Image size, 240 × 96 pixels$^2$ (acquired in HS-AFM observations); scan area, 80 × 64 nm$^2$; frame rate, 3.3 fps.

**Supplementary Video 4. HS-AFM videos of three representative CaMKIIα/GFP-CaMKIIβ 3:1 heterooligomers on a P[5]A+-modified mica surface.** The trajectories of the kinase domains are shown on the right. Green arrowheads and arrows indicate the kinase domains of CaMKIIβ subunits and GFP, respectively. Image size, 120 × 96 pixels$^2$ (acquired in HS-AFM observations); scan area, 80 × 64 nm$^2$ or 100 × 80 nm$^2$; frame rate, 3.3 fps.

**Supplementary Video 5. HS-AFM videos of three representative CaMKIIβ homooligomers treated with Ca$^{2+}$/CaM (1 mM Ca$^{2+}$ and 800 nM CaM) on a P[5]A+-modified mica surface.** The trajectories of the kinase domains are shown on the right. White dotted circles indicate Ca$^{2+}$/CaM bound to the kinase domains, and blue dotted squares indicate kinase domain complexes. Image size, 360 × 96 pixels$^2$ (acquired in HS-AFM observations); scan area, 80 × 82 or 80 nm$^2$; frame rate, 3.3 fps.

**Supplementary Video 6. HS-AFM videos of three representative CaMKIIα/β 3:1 heterooligomers treated with Ca$^{2+}$/CaM (1 mM Ca$^{2+}$ and 800 nM CaM) on a P[5]A+-modified mica surface.** The trajectories of the kinase domains are shown on the right. Green arrowheads indicate the kinase domains of CaMKIIβ subunits. White dotted circles indicate Ca$^{2+}$/CaM bound to the kinase domains, and blue dotted squares indicate kinase domain complexes. Image size, 240 × 96 pixels$^2$ (acquired in HS-AFM observations); scan area, 80 × 64 nm$^2$; frame rate, 3.3 fps.

**Supplementary Video 7. HS-AFM videos of three representative CaMKIIβ homooligomers treated with $Ca^{2+}$/CaM and ATP (1 mM $Ca^{2+}$, 800 nM CaM, and 1 mM ATP) on a P[5]A+-modified mica surface.** The trajectories of the kinase domains are shown on the right. White dotted circles indicate $Ca^{2+}$/CaM bound to the kinase domains, and blue dotted squares indicate kinase domain complexes. Image size, 360 × 96 pixels$^2$ (acquired in HS-AFM observations); scan area, 80 × 77, 72 or 76 nm$^2$; frame rate, 3.3 fps.

**Supplementary Video 8. HS-AFM videos of three representative CaMKIIα/β 3:1 heterooligomers treated with $Ca^{2+}$/CaM and ATP (1 mM $Ca^{2+}$, 800 nM CaM, and 1 mM ATP) on a P[5]A+-modified mica surface.** The trajectories of the kinase domains are shown on the right. Green arrowheads indicate the kinase domains of CaMKIIβ subunits. White dotted circles indicate $Ca^{2+}$/CaM bound to the kinase domains, and blue dotted squares indicate kinase domain complexes. Image size, 240 × 96 pixels$^2$ (acquired in HS-AFM observations); scan area, 80 × 64 or 100 × 80 nm$^2$; frame rate, 3.3 fps.

**Supplementary Video 9. HS-AFM videos of three representative CaMKIIβ homooligomers treated with EGTA (2 mM) and ATP (1 mM) after $Ca^{2+}$/CaM/ATP stimulation (1 mM $Ca^{2+}$, 800 nM CaM, and 1 mM ATP) on a P[5]A+-modified mica surface.** The trajectories of the kinase domains are shown on the right. Blue dotted squares indicate kinase domain complexes. Image size, 360 × 96 or 80 × 73 pixels$^2$ (acquired in HS-AFM observations); scan area, 80 × 74 or 73 nm$^2$; frame rate, 3.3 fps.

**Supplementary Video 10. HS-AFM videos of three representative CaMKIIα/β 3:1 heterooligomers treated with EGTA (2 mM) and ATP (1 mM) after $Ca^{2+}$/CaM/ATP stimulation (1 mM $Ca^{2+}$, 800 nM CaM, and 1 mM ATP) on a P[5]A+-modified mica surface.** The trajectories of the kinase domains are shown on the right. Green arrowheads indicate the kinase domains of CaMKIIβ subunits. Blue dotted squares indicate kinase domain complexes. Image size, 240 × 96 or 120 × 96 pixels$^2$ (acquired in HS-AFM observations); scan area, 80 × 64 nm$^2$; frame rate, 3.3 fps.

**Supplementary Video 11. HS-AFM videos of CaMKIIα$_{T286A}$ and CaMKIIβ$_{T287A}$ homooligomers treated with $Ca^{2+}$/CaM (1 mM $Ca^{2+}$ and 800 nM CaM) on a P[5]A+-modified mica surface.** An enlarged view of the white dotted box is presented on the right. White dotted circles indicate $Ca^{2+}$/CaM bound to the kinase domains, and white dots indicate the center of the kinase domains. Frame rate, 3.3 fps.
